# Analysis of the principles governing the dimensionality reduction performed by Lobula Plate Tangential Cells

**DOI:** 10.64898/2026.01.11.698863

**Authors:** Alix Leroy, Graham K. Taylor

## Abstract

Lobula Plate Tangential Cells (LPTCs) are critical for visual navigation in flies, enabling efficient processing of optical flow. Unlike upstream neurons in the visual pathway, such as those in the Lobula, Medulla, and T4/T5 layers, which retain a retinotopic organisation, LPTCs perform the first major dimensionality reduction of visual signals. This reduction simplifies the vast amount of visual information into key motion signals essential for navigation. In this paper, we investigate the principles governing this dimensionality reduction and explore with a data-based approach the efficiency of LPTCs in compressing optical flow information while preserving critical motion cues for navigation and their role in transmitting information to descending neurons. More specifically, our contribution is as follows: (i) we demonstrate, using Principal Component Analysis (PCA) as a reference, that global optical flow patterns, as also found in LPTC response maps, are efficient to describe and encode the variation in the optical flow experienced during visual navigation tasks; (ii) we show that LPTCs present a strong alignment with Principal Components of the experienced optical flow, supporting the hypothesis that the LPTCs maximize the flow of signal energy from the T4/T5 cells to the descending neurons; and (iii) we show that whilst LPTCs encode optical flow information in a lossy fashion, it can be reconstructed. Using optical flow priors, such as the complementarity of LPTCs in contralateral hemispheres and continuity of the flow in navigation tasks, we performed a reconstruction of the original flow from LPTCs’ response. Our analysis reveals the dual potential of LPTCs: selectively relaying signals for navigation that are fine-tuned in relation to the fly’s own dynamics; and broadly compressing the optical flow information that is sensed by the fly’s visual system.

## 1 Introduction

The visual navigation task, which involves estimating ego-motion in relation to the structure of the environment based on visual cues, has been a core focus of the computer vision community for over half a century. Classical approaches rely on accurately estimating motion through the detection of multiple points in the field of view representing key features of the scene. This is achieved using sparse optical flow algorithms [54], dense optical flow algorithms [30], or local feature-matching methods between successive image frames [53, 5, 11, 62]. These detected features are then used to extract relative pose information based on camera parameters [52, 56], often with the support of RANSAC algorithms to filter outliers in the selected key points [20].

Recent advances in deep neural networks have significantly improved these methods, enabling them to manage a wide range of scenarios, including indoor and outdoor environments, varying lighting conditions, large displacements, and sensor noise. However, despite their notable performance, these methods face two significant challenges. Firstly, state-of-the-art systems rely on deep networks for both optical flow estimation [69, 76] and local feature matching [13, 64, 66]. These networks often suffer from the so-called black box problem [26], which limits their explainability, interpretability, and the trust that can be placed in their predictions. This lack of transparency complicates the deployment of such systems in real-world scenarios. For instance, an unmanned aerial vehicle (UAV) is typically authorised to operate only once its systems have been thoroughly tested and certified by the relevant aviation authority. However, the “black box” nature of many machine learning systems makes it difficult to guarantee that the underlying algorithms have been evaluated across the full range of potential operational conditions. This issue becomes even more pronounced as artificial neural networks increase in complexity, rendering them susceptible to the curse of dimensionality [6], wherein the number of available training and testing data points becomes disproportionately small relative to the number of trainable parameters.

Secondly, to address the limitations of earlier approaches, modern deep networks have evolved to ensure accuracy across a broad range of scenarios. This has led to a dramatic increase in the number of trainable parameters, with some models now containing billions of parameters [14]. Such complexity imposes significant constraints on systems that rely on real-time information for navigation, including the need for substantial computational resources and energy.

In stark contrast to these modern computational methods, flies such as *Calliphora* and *Drosophila* demonstrate remarkable visual navigation capabilities using a biological system that is uniquely well understood and highly efficient [7, 68]. With a brain containing only ∼100,000 neurons and weighing just a few milligrams [77], flies have developed the ability to navigate complex environments over millions of years [40]. Their visual system provides a promising source of inspiration for the computer vision community, offering potential solutions for achieving explainable, accurate, and real-time navigation.

At the heart of the fly’s navigation system are the Lobula Plate Tangential Cells (LPTCs), a group of motion-sensitive neurons located in the visual system [42]. These neurons respond to optical flow, which is critical for insects performing tasks such as obstacle avoidance, course stabilisation, and orientation during flight [18]. Other pathways were found to be specifically tuned to looming stimuli [65, 39], although some studies suggest some rudimentary distance estimation capabilities of certain HS-cells [50] and VT-cells [51].

While the response profiles of LPTCs have been extensively characterised, the signal processing mechanism by which they compress visual information for self-motion estimation remain largely unexplored.

In the visual system of flies, earlier stages, such as the Lamina, Medulla, and the bushy T4/T5 cells, maintain a retinotopic organisation [7]. At these stages, the spatial arrangement of visual information is preserved, and the dimensionality of the input captured by the ommatidia remains proportional to the ommatidial array resolution *N*. However, after the local motion estimation by T4 and T5 cells in the four cardinal directions, these signals are relayed to LPTCs. Ouput LPTCs—commonly classified into Vertical System (VS) and Horizontal System (HS) cells—respond to vertical and horizontal optical flow patterns, respectively. These neurons are the first stage in the visual system where significant dimensionality reduction occurs. By integrating signals from multiple columns of the retinotopic array and providing single responses, LPTCs perform what we could describe as a compression of the visual information into structured outputs that are sent to descending neurons themselves providing navigation-relevant control signals [29, 72, 73]. Despite this known role, the exact principles underlying this dimensionality reduction and its contributions to visual processing remain key areas of investigation.

This biological dimensionality reduction allows LPTCs to compress signals while retaining critical structural and motion information thanks to LPTCs being less affected by fluctuations in the spatial structure of visual input compared to their presynaptic elements [12, 72]. This property allows neighbouring LPTCs to work as a compression system robust to noise. Similarly, Principal Component Analysis (PCA) [31], a well-established technique for dimensionality reduction, offers a computational counterpart. PCA is frequently used to simplify complex datasets by projecting them onto orthogonal components that capture the most significant variations. Notably, PCA has been shown to enable control in a lower-dimensional space while managing structural instabilities in dynamic systems [55], making it a comparable framework for understanding the data compression performed by LPTCs.

In the past, PCA has been widely applied in the context of optical flow to enhance estimation and performance in various scenarios, yet never with the goal to solve ego-motion through dimensionality reduction. For example, PCA was successfully used in optical flow estimation using an event-based camera, known as neuromorphic cameras [59], where traditional frame-based approaches may struggle. The design of these sensors is based on biological vision with sparse event-based output, a representation of relative luminance change, and its encoding of positive and negative signals into separate output channels [49]. In the fly’s visual system, such a functional phenomenon can be observed in the Lamina with the encoding of brightness change into the ON and OFF pathways [35, 36]. Neuropils founds further down in the visual system, such as the Medulla, or the Lobula hosting the inputs of T4/T5 cells provide functional capabilities of higher level than the event-based cameras, such as direction selectivity. PCA was successfully used with event-based cameras optical flow estimation [38], incorporating regularisation techniques to enhance performance. This approach, designed for real-time visual odometry, achieved a twofold speed improvement over state-of-the-art methods while significantly enhancing optical flow accuracy, thus demonstrating the usefulness of PCA in scenarios requiring efficient and accurate processing, such as real-time navigation systems.

In addition to enhancing optical flow estimation, PCA has been explored to analyse the natural components of optical flow fields. PCA-Flow, introduced by Wulff et al. [75], learns the principal components of natural flow fields to efficiently compute dense optical flow. By assuming that optical flow lies on a low-dimensional subspace, PCA-Flow enables fast computation of approximate optical flow. This approach was further extended with PCA-Layers[75], which incorporated a layered model to improve accuracy, particularly at boundaries. Despite these advances, these methods focus on enhancing existing optical flow estimation rather than encoding optical flow accurately or uncovering its intrinsic structure.

Efforts to analyse the natural components of optical flow have also highlighted the importance of distinguishing between rigid and non-rigid motion components. For example, Fleet et al. [22] proposed linear motion models to effectively describe optical flow, using “motion features” such as motion discontinuities and moving bars, as well as object-specific motions. These linear parametrised models provide strong constraints on the spatial variation of optical flow and offer a concise description of motion through a small set of linear coefficients. This is analogous to the fly’s visual system, where dimensionality reduction is achieved while retaining essential motion cues. The model coefficients can be directly estimated from image derivatives, bypassing the need for dense motion computation.

For motion features, Fleet et al. [21] constructed a basis of “steerable flow fields” to approximate motion features like occlusion boundaries and facial expressions. Similarly, object-specific motions were analysed using principal component analysis to construct basis flow fields from example motions with no prior computation of optical flow. These coefficients were shown to be useful for detecting and recognising specific motions, further validating the importance of PCA in describing and analysing of perceived motion.

Building on earlier research, Runia et al. [63] extended the decomposition of flow fields to include broader motion patterns using the Divergence-Curl-Shear (DCS) descriptor [34]. This work continued the trajectory established by foundational studies such as the ones of Koenderink and Van Doorn [41], and Abraham et al. [60], highlighting the utility of global optical flow patterns as more than just tools for dimensionality reduction. Instead, they offer insights into the underlying structure of optical flow fields, providing a framework to analyse and interpret motion in dynamic environments.

The DCS descriptor effectively breaks down optical flow into divergence, curl, and shear components, capturing specific motion patterns in the field of view. It differs from the methodology employed in this study, where PCA is used to extract motion patterns with components specifically tuned to the dynamics of the agent, as they are defined by the maximisation of variance in the data rather than by the mathematical structure of the motion Jacobian on which the divergence, the curl and the shear components rely.

Moreover, the PCA-derived components employed in this study are applied globally to the entire field of view, rather than being derived from filter designs that convolute over the flow map, as in the DCS approach. This global perspective allows the principal components to capture dominant patterns across the visual field, reflecting the broader motion cues experienced by the agent. By contrast, the DCS descriptor provides a more localised analysis of motion patterns, which may be beneficial in certain scenarios, such as looming detection, but does not account for the global dynamics.

By examining the role of LPTCs in dimensionality reduction and comparing it to PCA on optical flow data, this study seeks to bridge biological and computational principles of visual navigation. Insights derived from studying LPTCs have the potential to inspire innovative solutions in computer vision, particularly for applications requiring real-time, efficient, and explainable navigation systems.

Specifically, our contributions are as follows: (i) We demonstrate that global optical flow patterns, similar to those observed in LPTC response maps, effectively capture and encode the variance in optical flow data experienced during navigation tasks, as revealed through PCA. (ii) We show that LPTCs exhibit strong alignment with the principal components of experienced optical flow, suggesting their role in maximizing the transfer of signal energy from T4/T5 cells to descending neurons. (iii) We demonstrate that while LPTCs encode optical flow information in a lossy manner, optical flow priors, such as the complementarity of LPTCs in the contralateral hemispheres and the continuity of flow during navigation, enable the reconstruction of the original optical flow from LPTC responses.

The subsequent sections will detail our methodology, present the results of our comparative analysis, and discuss the broader implications of these findings for both biological and computational models of visual processing.

## 2 Methods

### 2.1 Data Processing

The FlyView dataset [47] is a bio-informed optical flow dataset developed for visual navigation tasks, leveraging panoramic stereo vision to mimic the visual system of insects, particularly *Calliphora* and *Drosophila* flies. It was designed to facilitate the exploration of the fly visual system and address challenges faced by robotic and autonomous systems in navigating complex environments. FlyView incorporates several key features such as panoramic images with a wide-angle field of view resembling that of a fly’s compound eyes. Additionally, the dataset is based on motion primitives and realistic 3D trajectories of drones and flies, simulating dynamically-meaningful navigation scenarios, and provides pose and motion flow ground truth for each time-step.

For this study, we utilised over 7,200 instances of fly trajectories from FlyView (subsets *S*7, *S*8, and *S*9), as well as 5,387 instances of its drone trajectories in a separate dataset used for comparison (free flight trajectory *S*5_0_). The input data consisted of motion flow maps, representing the groundtruth optical flow experienced by the agent during various self-motion scenarios. These maps were generated using FlyView’s Blender simulation, which ensures pixel-accurate representations of optical flow without the limitations commonly associated with computer vision-based optical flow estimators, such as noise [70, 27], occlusion [33, 3], illumination variations [71, 58, 2], or large displacements [8, 9].

The motion flow maps were resized and cropped from FlyView’s original high-resolution dimensions (1700 × 900px) to match the 25 × 11px resolution of LPTC response maps [42, 44, 45, 43, 46]. Some experiments, discussed later, used larger input resolutions to evaluate the effect of resolution on explained variance. We found that the optical flow maps were very sensitive to the method used to resize the flow map. In total, we tested five different downsampling methods including bilinear interpolation, bi-cubic interpolation, nearest neighbour interpolation, Lanczos interpolation, and a local averaging. Whilst the first four methods led to distorted flow maps, the local average resulted in consistent resized flow maps which could also be reconstructed back to their original size with without artefacts. The LPTC responses to these flow maps were then estimated using a circular statistics model [25, 48], which describes the tuning properties of individual cells. Each cell is selectively tuned to specific aspects of optical flow, including distinct directions and magnitudes of rotational and translational motion. The LPTC dataset comprised a total of 26 cells: 10 VS cells and 3 HS cells for each hemisphere.

To enforce symmetry across the dataset, we applied a data augmentation technique by mirroring each optical flow map along the 0-degree azimuth axis, i.e. the thrust direction. This transformation was performed independently for both the fly and drone datasets, generating two fully symmetrical versions. By implementing this augmentation, we doubled the number of instances and we ensured a balanced representation of motion patterns, reducing potential directional biases in the analysis. Figures 1a and 1b shows an instance flow map from FlyView before and after data augmentation.

**Figure 1:**
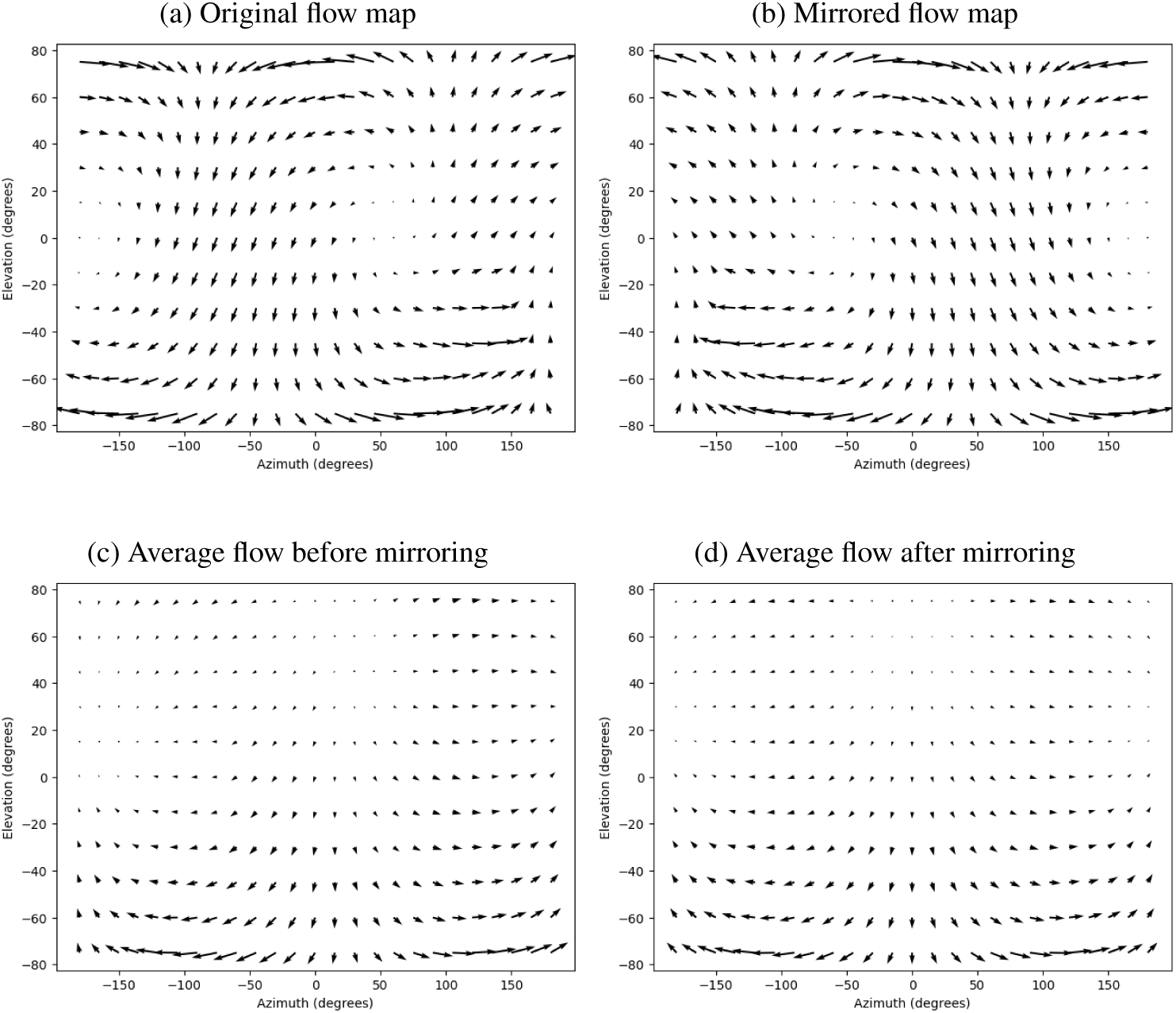
Data augmentation visualisation. (a) Motion flow instance from fly trajectories present in FlyView. (b) Example of mirrored Motion Flow used for data augmentation. Average Optical Flow experienced in the fly trajectories before (a) and after (b) data augmentation mirroring the optical flow maps. Without the mirroring of the optical flow maps, the dataset presents an asymmetrical forward motion pattern whose centre is located around 35 degrees azimuth. After performing the mirroring operation, the resulting average optical flow is perfectly symmetrical along the 0-azimuth axis.

Figure 1c shows the optical flow obtained when averaging the local vectors across the selected fly dataset. We can observe that it contains mostly a forward translation component on the ventral part of the field of view. This optical flow is slightly asymmetrical along the 0-azimuth axis. Mirroring each instance of the fly dataset resulted in a perfectly symmetrical average flow field as shown in Figure 1d.

### 2.2 Principal Component Analysis

Principal Component Analysis (PCA) was used in this study as a comparison to the Lobula Plate Tangential Cells (LPTCs). Unlike previous applications of PCA, which often focus for optical flow estimation or on specific regions or subsets of the flow field, our approach applies PCA to an already estimated optical flow, similarly to LPTCs, and to the entirety of the optical flow field. To the best of our knowledge, this is the first instance where PCA has been used with the explicit goal of analysing global optical flow patterns in this manner. The reason such an approach has not commonly been pursued may lie in the assumption that PCA will only yield interpretable principal components when applied to optical flow fields that are structured by a consistent source of motion, such as self-motion. In the case of rigid-body self-motion, the resulting optical flow fields exhibit coherent and predictable global patterns, which PCA can then decompose into meaningful components. By contrast, in scenarios involving multiple independently moving objects or highly variable scene geometry, PCA may struggle to extract consistent patterns. Therefore, the use of PCA as a tool for understanding global optical flow structure is particularly well-suited to the analysis of biologically relevant motion, such as the self-generated visual input experienced during free flight.

The PCA computation was conducted using Singular Value Decomposition [28], as described by Moore [55]. To ensure robust dimensionality reduction and remove bias, we performed a centred PCA, subtracting the average optical flow from each dataset prior to computation. This preprocessing step ensures that the PCA captures the variance in the data rather than the mean optical flow, leading to more accurate principal components.

Similarly to the original data processing, the input optical flow maps were resized to a resolution of 25 × 11 vectors by averaging neighbouring local vectors of the flow field, reflecting the dimensions of the LPTC response maps. This resizing resulted in 550 components being derived from the PCA. These components were then ranked by the variance they explained, allowing for a detailed analysis of the dimensionality reduction process and a comparison to the LPTC responses encoding efficiency.

PCA was applied to both the fly and drone datasets to explore differences in the optical flow fields that these flying agents experience as a result of their respective flight dynamics. Additionally, we tested scenarios that included the full field of view of the fly, as well as cases where the left and right hemispheres were masked at a −15 and +15 degrees azimuth angle, respectively. This separation aligns with the field of view observed for individual compound eyes, enabling a more representative comparison of hemispheric contributions to optical flow encoding.

The Explained Variance Ratio in PCA quantifies how much of the total variance in the data is captured by each principal component. It is computed using the eigenvalues obtained from the covariance matrix of the data.

The eigenvalues *λ_i_* of the covariance matrix represent the variance captured by each corresponding eigenvector (principal component). The total variance in the data is the sum of all eigenvalues.

The explained variance ratio for each principal component *i* is given by the ratio of its eigenvalue *λ_i_* to the sum of all eigenvalues. This ratio indicates the proportion of the dataset’s total variance that is captured by the *i*-th principal component.

Mathematically, the explained variance ratio EVR*_i_* for the i-th principal component is given by:

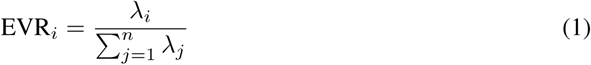

where:

- *λ_i_* is the eigenvalue corresponding to the *i*-th principal component.
- *n* is the total number of principal components (or the total number of features in the original data).

Often, we are also interested in the cumulative explained variance ratio, which is the sum of the explained variance ratios of the first k principal components. It tells us how much of the total variance is captured by the first k principal components:

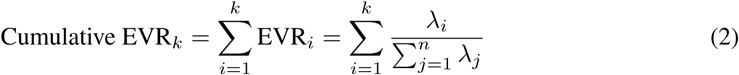

The cumulative explained variance ratio was estimated on the datasets at different resolutions (25×11, 50 × 22, 100 × 45, and 250 × 110 pixels) to highlight the impact of input resolution on the distribution of energy in the system.

### 2.3 Decoding the LPTC response

LPTCs encode optical flow-based information in global patterns, akin to Principal PCA, but they differ in several significant ways. First, the LPTC model of Leroy et al. [48] based on the linear model defined by Krapp et Hengstenberg. [45] to model the response before the plateau of saturation [25], encodes only positive values along the Local Preferred Direction (LPD). This leads to a suppressed response along the Local Null Direction (LND). This creates a limitation in encoding local motion patterns as the simulated response of our model will yield outputs on floating points numbers that are only positive, oppositely to the PCs that will give positive and negative values when applied due to their linear encoding. Secondly, to compensate this first limitation impacting the ability to discriminate some motions from one another, such as pitch rotation and lift translation, LPTCs are divided between two hemispheres, with the response maps of one hemisphere responding oppositely along the azimuth axis to those of their contralateral counterparts. Thirdly, LPTCs do not cover the entire field of view of their own hemisphere, thus remaining unresponsive to some regions of the field of view. As a result, stimuli along the LND, in the opposite hemisphere, or outside the response field of a cell can all lead to non-responsiveness in numerous LPTCs.

To process visual information efficiently despite these constraints, it is likely that the LPTCs rely on a combination of multiple cells, if not all, to describe the experienced optical flow locally. We test whether the LPTCs output can be used to reconstruct the optical flow experienced by the fly. However, the severe limitations of PCA in decoding optical flow data (See results in 3.3) highlight the need for alternative approaches for efficient decoding. Based on the limitations and specific characteristics of the LPTCs’ lossy compression of information, described above, we propose that a non-parametric or non-linear method would be better suited for this task. Given that both LPTC responses and optical flow maps can be described, when rescaled, as distributions, we opted for a Gaussian Process (GP) Regression model [61] in this study.

Gaussian Processes are a non-parametric supervised learning method used to solve regression and probabilistic classification problems. This approach offers several advantages. First, GP predictions interpolate the observations, at least when using regular kernels. Second, predictions are probabilistic (Gaussian), allowing for the computation of empirical confidence intervals. These intervals can be leveraged to decide whether additional data is needed for model refinement in certain regions of interest. Finally, Gaussian Processes are versatile, as different kernels can be specified depending on the data. Common kernels are readily available, but custom kernels can also be defined, making the method highly adaptable to specific challenges.

In our case, we applied GP Regression to both fly and drone trajectories. Since Fly and drone trajectories have different magnitude of optical flow vectors, thus leading to different magnitude of response for LPTCs, we decided to normalize both datasets by the maximum value of the latent space, i.e the value of the cell responding maximally to the optical flow map. To ensure the results remained unaffected by this normalization, and because LPTC responses provide a linear relationship to the magnitude of the flow experienced, we reintroduced the magnitude of the flow at the final stage when computing these metrics, more specifically for the End-Point Error (See Section 2.4 for a more detailed description of these metrics).

The kernel used for our GP model was the Squared Exponential (SE) kernel, defined as:

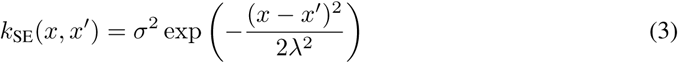

The Gaussian kernel is the *de facto* standard for Gaussian Processes and Support Vector Machines (SVMs). This popularity stems from its universal applicability and desirable mathematical properties. Every function within its prior has infinitely many derivatives, making it highly smooth and versatile. The kernel is governed by two parameters: the lengthscale *λ* and the output variance *σ*^2^. The lengthscale determines the scale of “wiggles” in the function; generally, the model cannot extrapolate beyond a distance of *λ* from the data. For our specific case, we fixed the lengthscale due to memory requirements, which exceeded 300 GB of RAM to initialize the model. This constraint also precluded the use of SVMs for this challenge. The output variance *σ*^2^, on the other hand, determines the distance of the function from its mean value, acting as a scale factor.

### 2.4 Optical Flow Metrics

In order to provide an experimental setup where optical flow estimation algorithms can be reasonably tested and compared, a group of metrics has been defined as standard. The first papers dealing with direct quantitative evaluation metrics for optical flow were published in 1994 and suggested two main metrics. The first, the End-Point Error (EPE) [57], can be described as the Euclidean distance between two vectors *G* = (*u, v*) and *E* = (*u^∗^, v^∗^*); it is defined in Equation 4.

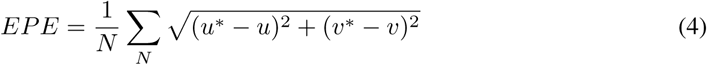

The second is named the Average Angular Error (AAE) ([4] based on prior work of [23]) which represents the angle between the two extended vectors *G_e_* = (1*, u, v*) and *E_e_*= (1*, u*\**, v*\*) and defined in Equation 5:

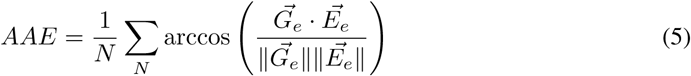

AAE is highly sensitive to small estimation errors caused by small displacements, whereas EPE hardly discriminates between close motion vectors [24].

Even though EPE and AAE metrics are popular, neither metric is better than the other and, therefore, they can be considered complementary. For example, AAE penalizes errors in regions of zero motion more than motion in smooth non-zero regions. In addition, there are different cases where EPE outputs the same value in very different scenarios [1]. This is caused by the fact that EPE considers only the difference of vectors and ignores the original magnitude of each one (See Figure 2).

**Figure 2:**
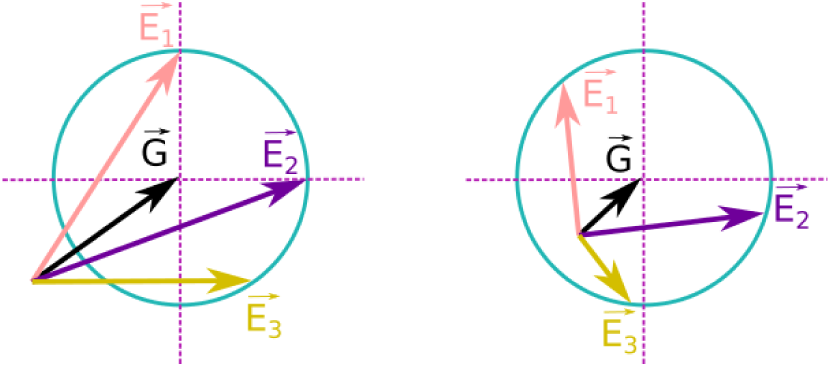
Representation of the End-Point Error (EPE). Two scenarios in which the EPE metric yields identical error values between the ground truth, represented by the black vector G, and various estimated optical flow vectors. Let E_1_, E_2_, and E_3_ denote three distinct estimations. As the EPE metric focuses solely on the distance between the endpoints of the estimated vectors and the ground truth, both cases (left and right) result in equivalent error values. (Adapted from [1]).

These metrics are being used in this paper both for the PCA analysis, through reconstruction of the original signal from the inverse Principal Components usage, and in the Optical Flow decoding based on the Gaussian Process Regression model.

## 3 Results

### 3.1 Qualitative Assessment of Principal Components

We observed a striking similarity between the principal components obtained after fitting the PCA on fly trajectories and the response maps of LPTCs. Most notably, the first Principal Components computed on the entire field of view describe coherent patterns similar to the LPTCs response maps (See Figure 4). Oppositely to LPTCs which couple rotation and translation and whose response fields have systematic variation in sensitivity across the visual field, these coherent patterns found in the first Principal Components describe optical flow templates covering the entire field of view and seem to separate translations and rotations motions whose axes are close to the principal axes of the flying agent. Notably, we identified motions such as forward translation (PC7), yaw rotation (PC1), pitch rotations (PC2), and lift down (PC4). Additionally, PC3 is mostly roll sensitive with a small elevation offset for its rotation axis, involving some yaw component in the pattern. Components of lesser importance appeared increasingly chaotic and incoherent (See 15 in Supplementary Material).

The results remained consistent when using both centred and uncentred PCA, i.e without subtracting the average optical flow from the dataset prior to computation. The principal components in the uncentered PCA closely mirrored those obtained from the centred version. For components that differed, changes were limited to their ranking or the appearance of opposite patterns. This suggests that centring the data, thus processing the optical flow represented as a deviation from the mean optical flow field experienced in forward translation flight, does not fundamentally alter the patterns captured by PCA but may influence their relative importance.

When the PCA was fitted on a mirrored dataset, we observed a symmetry in the principal components for the left and right hemispheres, whereas the original, non-mirrored dataset displayed dissimilar components highlighting the asymmetry in the FlyView data prior to data augmentation.

Finally, we found that the Principal Components derived from blowfly and quadcopter trajectories within FlyView present similarities; however, they appeared to capture self-motion components differently. Indeed, PC1 on drone data also responds to yaw rotations but does it in such a way that the local preferred directions of the vectors are almost perfectly horizontal. Additionally, we observed that PC2 and PC3 from drone data both encode pitch rotations by dividing the field of view in a upper and lower part. Similarly, PC4 and PC5 respond to roll sensitive motions dividing again the field of view in a upper and lower part. Finally, PC6 responds to lift motions and lower rank PCs do not correspond to any specific motions.

Masking one hemisphere for fly trajectories resulted in principal components whose patterns closely matched the ones found when processing the entire field of view (See Figure 3). However, we found that masking one hemisphere for drone trajectories resulted in patterns varying from their full field of view equivalent from the second component. Particularly, splitting the experienced optical flow into different hemispheres led to the first three PCs of both fly and drone trajectories aligning strongly. Additional visualisations of the Principal Components are available in Supplementary Materials.

**Figure 3:**
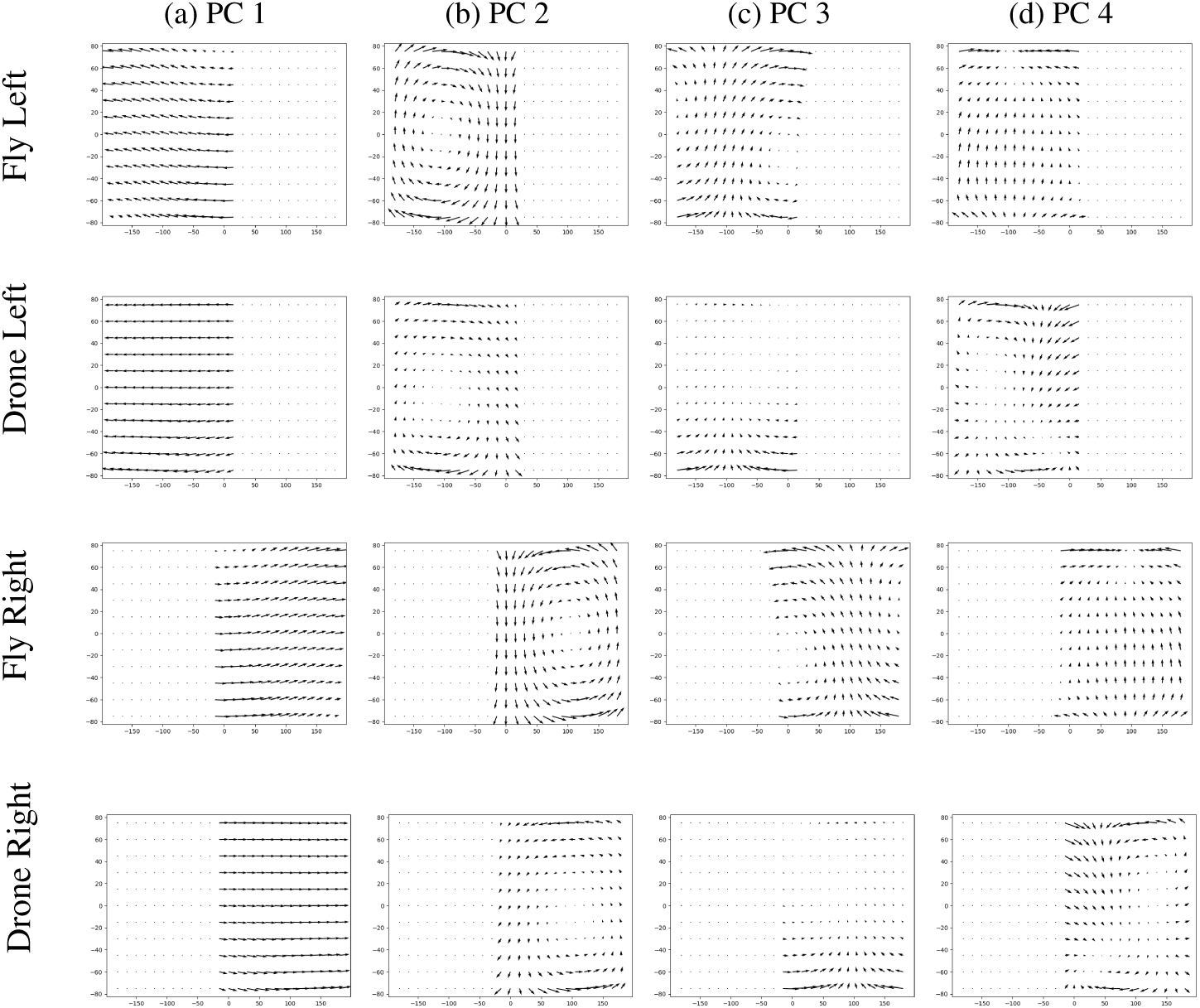
First four Principal Components (PC) computed on fly and drone datasets masking the Right and Left part of the field of view. PCs mimic the field of view of each eye in *Calliphora*. Rows correspond to, top to bottom: (1) Left hemisphere on fly trajectories, (2) Left hemisphere on drone trajectories, (3) Right hemisphere on fly trajectories, and (4) Right hemisphere on drone trajectories. When fitting the PCA on the augmented dataset (with mirrored optical flow maps), the resulting Principal Components on the left and the right hemispheres end up mirrored.

**Figure 4:**
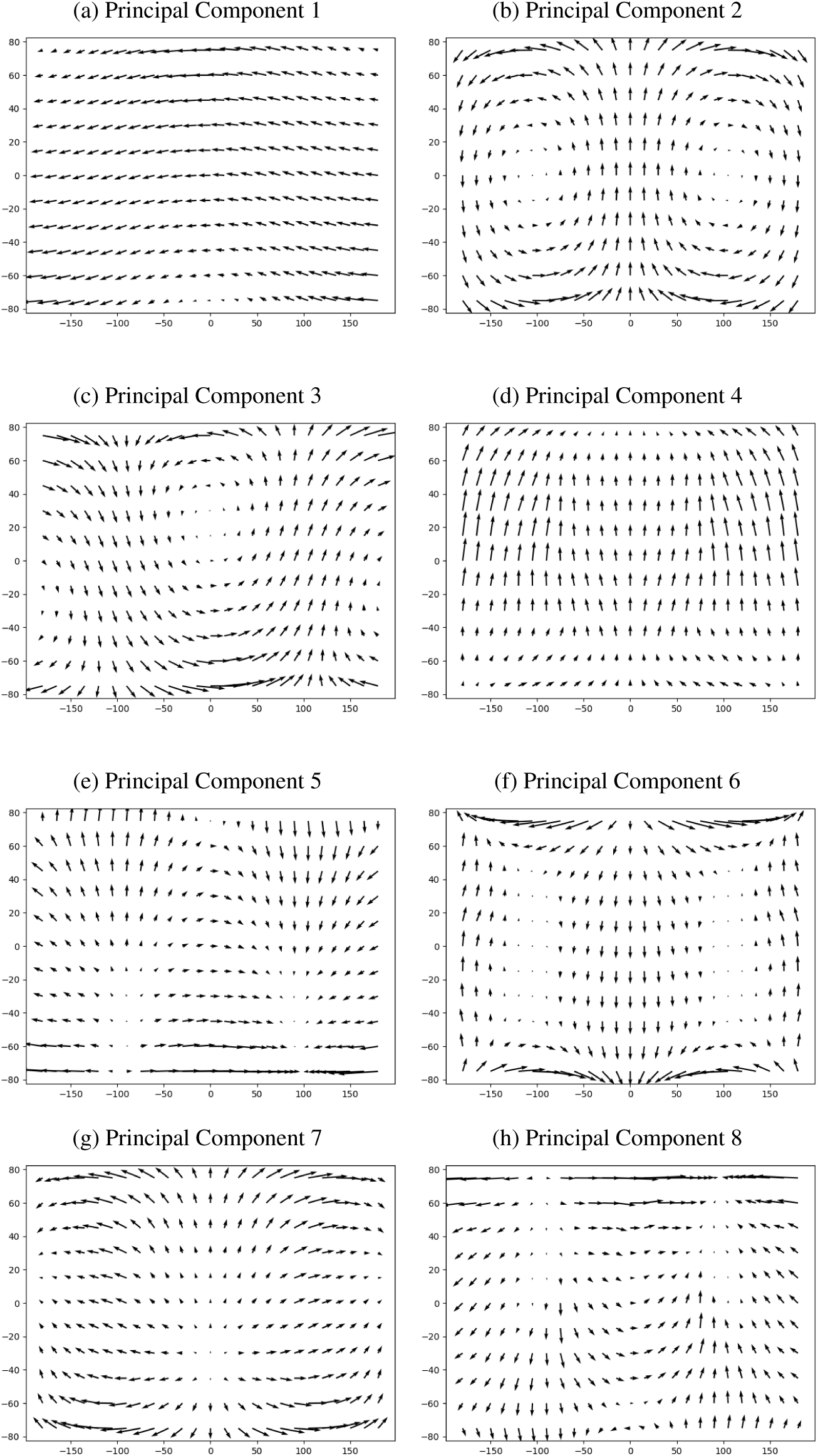
First Principal Components fitted to optical flow of fly trajectories from FlyView.

Whilst the LPTCs response map do not match entirely with the Principal Components we extracted, neither in the scenario of fly trajectories nor drone trajectories, LPTCs and PCs share numerous similarities in the coherent patterns, with VS and HS cells corresponding to rotational and translational patterns, respectively.

The analysis of the alignment of LPTCs’ preferred directions [25, 32] with PCs reveals a significant alignment of LPTCs with some PCs (See top row of Figure 5). Particularly, VS-cells were found to have a strong alignment with PC2 (VS1-VS3 and VS8-VS10) and PC3 (VS4-VS7) but also to align with PC1 with different response for each hemispheres corresponding to yaw rotations, and to align slightly with PC4 permitting lift down motion detection in the absence of response from both hemispheres. The HS-cells were found to strongly align with PC1 and PC7 highlighting their capabilities to detect forward and yaw rotation.

**Figure 5:**
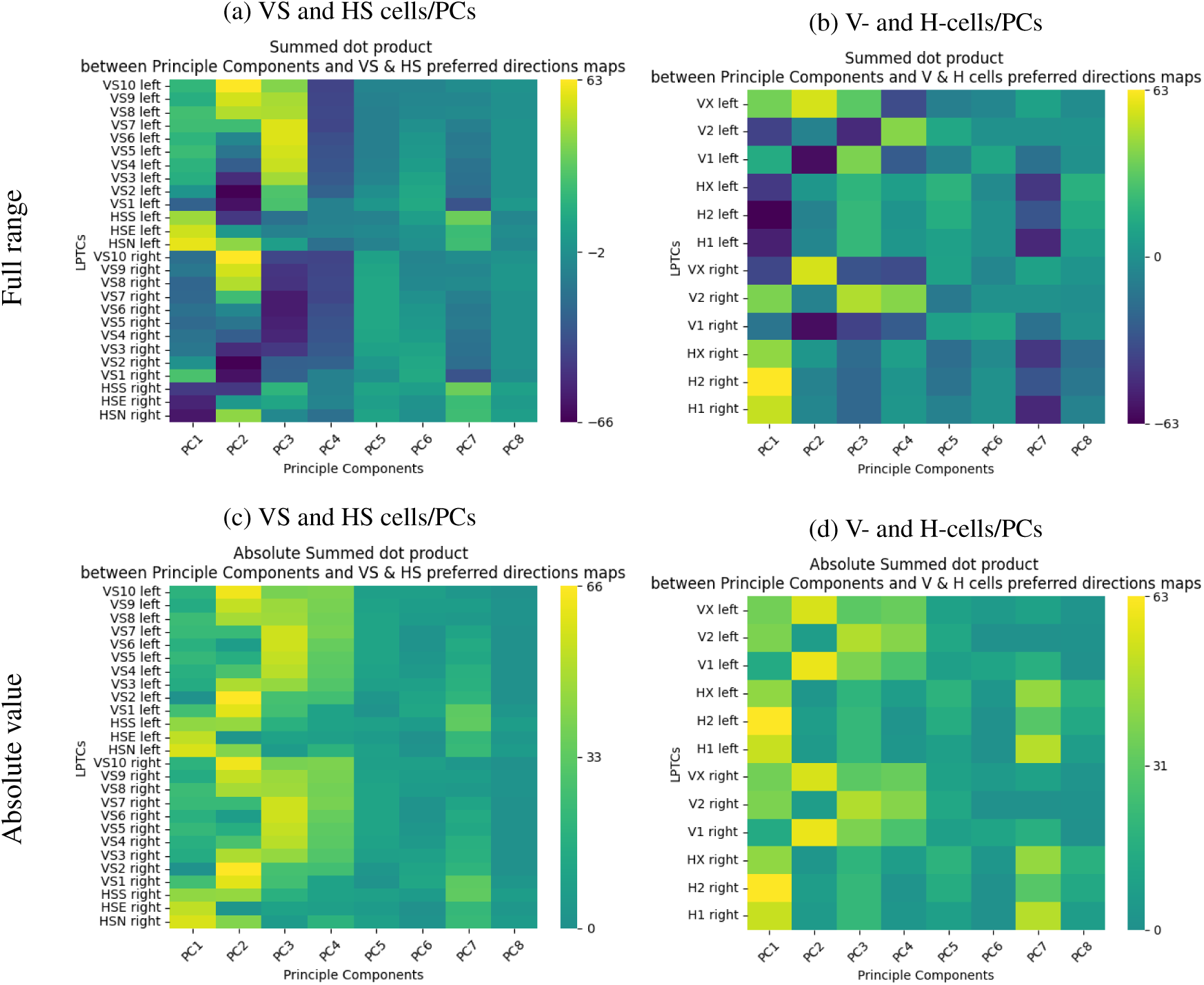
Alignment of the first PCs computed on fly trajectories from FlyView LPTCs include VS and HS cells (left) and V- and H- cells (right). Top row corresponds to the original alignment, allowing us to appreciate the differences between hemispheres Bottom row is the absolute value of the alignment, highlighting the close match between VS/HS responses with V/H responses.

Comparing the absolute alignment (absolute summed dot product) of the VS/HS cells group with the V/H cells group (See bottom row of Figure 5), we observe a similar structure between the heatmaps with V-cells responding similarly to VS-cells and H-cells responding similarly to HS-cells. This highlights the role of V/H cells to describe as an even more lower dimensionality and up to a sign, the optical flow experienced by the fly.

### 3.2 Analysing the Explained Variance

The resolution of the input data influenced the number of components required to capture variance. Higher resolutions demanded more components to explain the same proportion of variance. For instance, in all scenarios, we found that 26 components were sufficient to describe the variance of optical flow maps with 95% accuracy. Specifically, the unilateral hemisphere required 13 components, while the combined hemispheres required 15 components to achieve this level of accuracy (see Figure 6).

**Figure 6:**
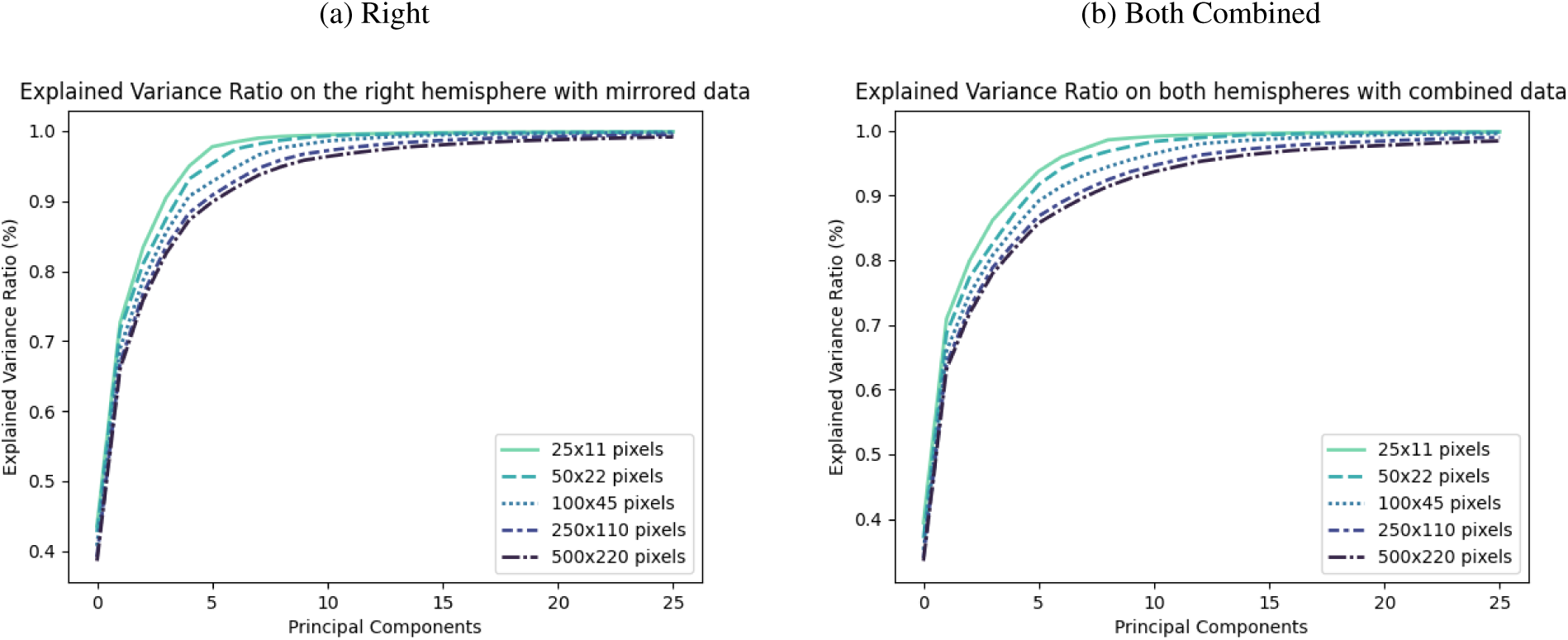
Evolution of Explained Variance Ratio. Comparison of the explained variance ratio with respect to the number of principal components used and the sensor resolution. Four scenarios are compared based on the data they experience. For the “Right” scenario (a), the flow maps are experienced only on the right hemisphere, thus reducing the number of photoreceptors experiencing optical flow at the same time, and providing Principal Components covering only the right hemisphere. This scenario is akin to ipsilateral cell spanning over a single hemisphere. For the “Both combined” scenario (b), the optical flow is experienced over the entire field of view collected by a monocular camera, providing Principal Components covering both hemispheres, similarly to heterolateral LPTCs.

Interestingly, increasing input resolution did not lead to a dramatic rise in the number of components required to explain variance beyond a certain threshold. The Left, Right, and Both Combined scenarios appeared to reach a limit in the loss of information. We hypothesize that this plateau occurs due to two main factors: (a) the static nature of the observed scenes and (b) the dynamically meaningful motions leading to specific optical flow experiences by the fly. These factors will constrain the variability of the optical flow maps, thereby limiting the number of principal components required to represent them. As a result, when considering the little contribution to explain the variance for lower rank Principal Components and the chaotic patterns present in them, it is very likely that such components are required solely for the consistency in the data representation with a linear approach rather than the description of its content that could be used for a navigational task.

### 3.3 Decoding Optical Flow

The particularity of the PCA is its linearity allowing to easily encode information and decode it back to its original form with little, if any, loss.

Processing the PCA on motion flow maps experienced during fly trajectories from FlyView, we were able to reconstruct the original optical flow fields using the entire set of principal components and applying the inverse operation. Figure 7 shows the variation of AAE across percentiles of the dataset for different resolutions (respectively 25 × 11px and 100 × 45px) for a range of components varying from 10 to the maximum number possible. The number of components (550, and 9000) varies with the resolution of the optical flow estimated (25 × 11*px*, and 100 × 45*px* respectively).

**Figure 7:**
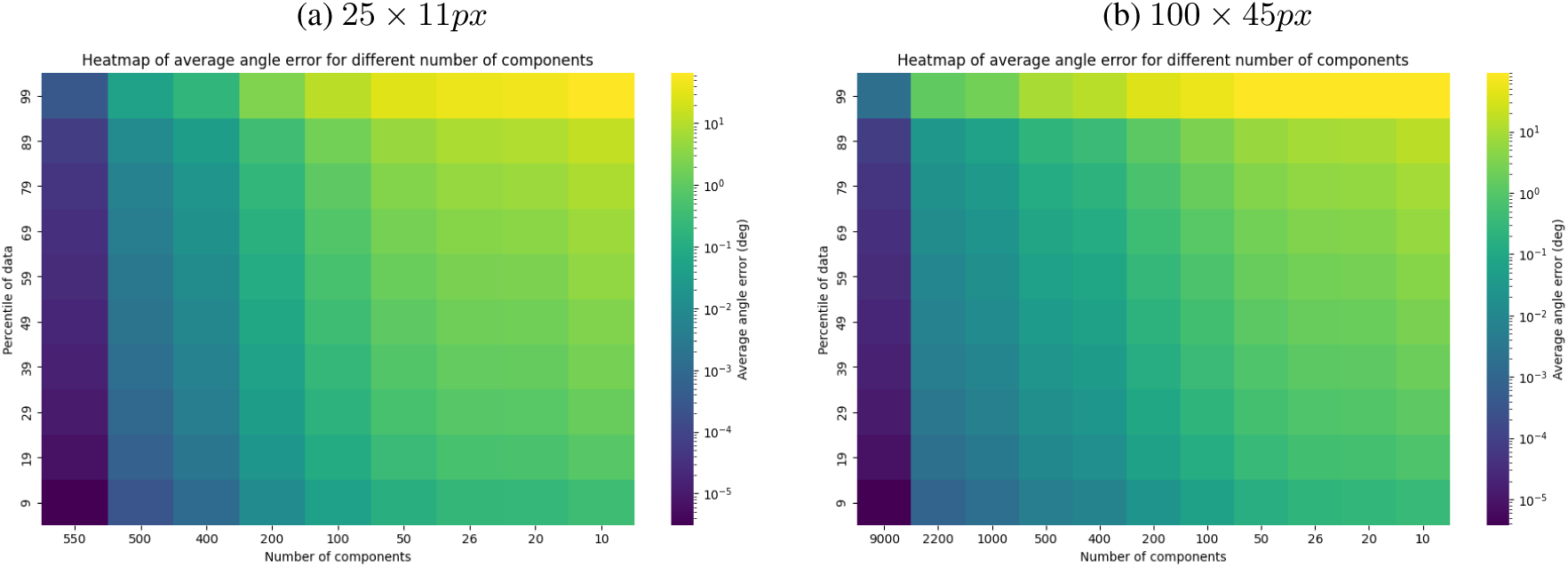
Heatmaps of error percentiles based on PCA-based linear decoder on the fly dataset. Percentiles of AAE computed (Y-axis) on individual optical flow from the fly dataset encoded by the *N* Principal components (X-axis) and decoded using the linear inverse function. The percentiles are shown for two different optical flow map resolutions: (a) 25 × 11*px* and (b) 100 × 45*px*. The percentiles of AAE are computed from the 9*^th^* percentile to the 99*^th^* percentile. We observe that whilst the linear encoding-decoding operation with the maximum number of components allows a reconstruction with a high accuracy on the AAE - as shown by the small errors for the 99*^th^* percentile of both resolutions, indicating as almost perfect reconstruction of the flow directions- the AAE increases as soon as we start removing some low-rank components. Whilst we observe that Principal Components have the potential to encode and decode the optical flow maps, it appears that the linear approach leads to exponentially increasing angular errors.

Whilst PCA captures well the variance of the dataset with a limited number of components, as soon as we reduced the number of components used in the reconstruction of the original signal, the accuracy of the optical flow generated, observed through the AAE, dropped dramatically thus excluding parametric linear approaches to offer an accurate way of encoding optical flow information in such a way it can be decoded. Considering the large amount of Principal Components, even for a small optical flow map encoding, the low rank principal components, containing chaotic patterns, are contributing to the encoding of the specific dataset for which they were fitted. When removing these components, they are not compensated in anyway by the linear operation. The low rank components, while encoding background noise of the dataset used to fit the data, ensure the accuracy of the signal reconstruction on this specific dataset.

In the case of tangential cells encoding optical flow patterns, their number is significantly limited within the Lobula Plate, with approximately 60 such cells in *Calliphora*. Given that *Calliphora* possesses around 10,000 ommatidia across its two compound eyes, the ratio of Lobula Plate Tangential Cells (LPTCs) to sensory input is approximately 1:166. By contrast, using an optical flow map of equivalent resolution (100×100 pixels) allows for the extraction of 20,000 Principal Components (PCs), providing 332 times more descriptors at an equivalent dimensionality of sensory input.

Furthermore, LPTCs are distributed across two separate hemispheres and do not span the entire field of view of the compound eye behind which they are located. These anatomical constraints suggest that LPTCs encode optical flow information in a manner that is less complete—and potentially more lossy—compared to a full representation of the optical flow field. Consequently, due to these inherent limitations, it is unlikely that the original optical flow can be reconstructed using a purely linear approach, as is possible with Principal Component Analysis (PCA).

However, the LPTCs having an organised distribution of the patterns, we can rely on self-motion and flow field priors to retrieve the original optical flow. Indeed, where the PCs show templates aligned with orthogonal directions, thus losing spatial coherence beyond PC8, LPTCs have the advantage to have both intra- and inter-hemispheric coherence in its template distribution. This distribution, inherited from the biological evolution of the fly’s visual system, supports the possibility to reconstruct the original optical flow with a different method. As a first attempt to test this hypothesis, we used the Gaussian Process Regressor which allowed us to reconstruct the optical flow maps (for 25×11px maps) with a high accuracy using LPTCs (See Figures 8a and 8b). We observe that estimations outputted by the model when being fed by the 26 LPTCs allows a fine reconstruction of the global flow. Some artefacts appear sporadically throughout the field of view, but the global consistency of the flow is preserved.

**Figure 8:**
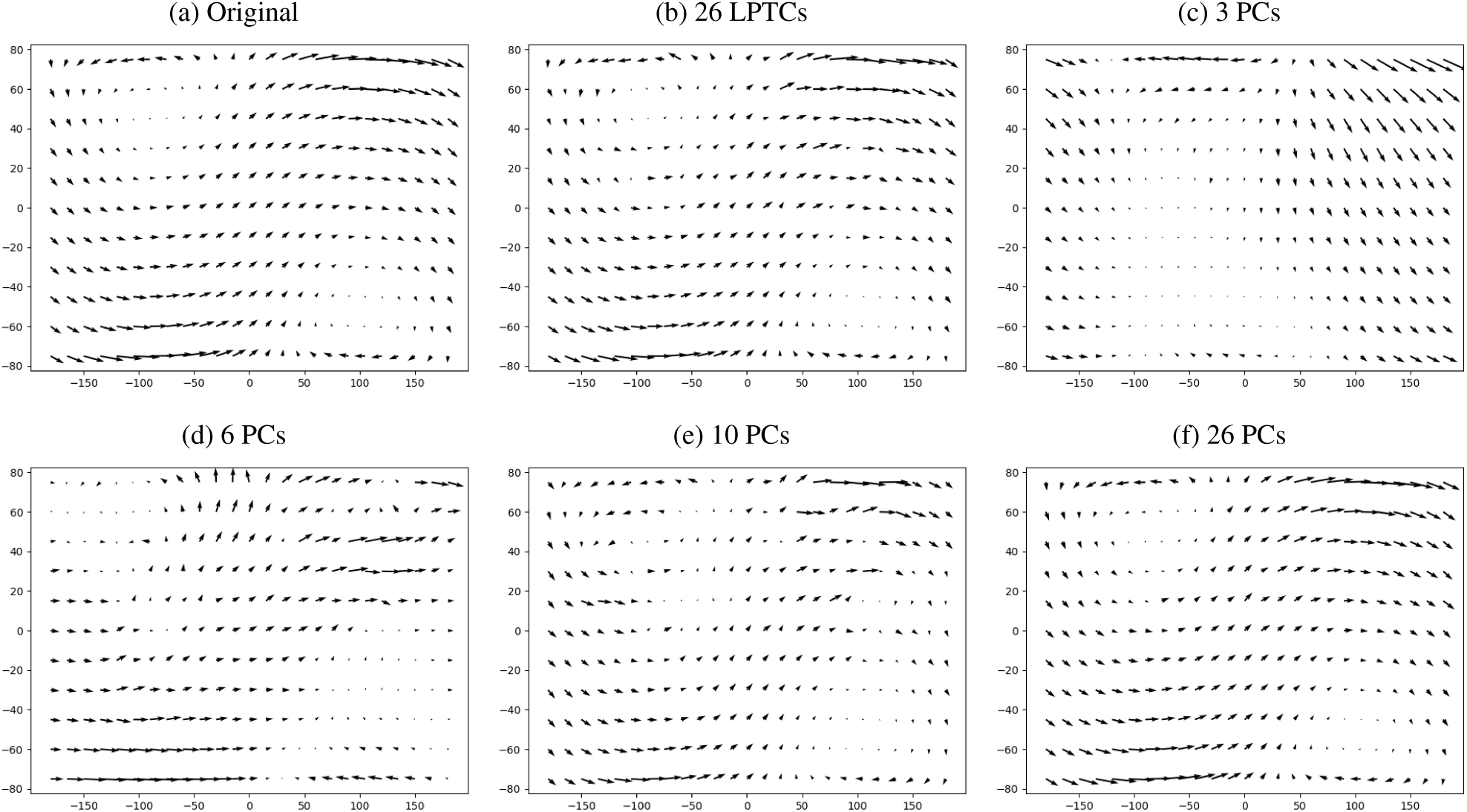
Comparison of decoded optical flow based on the encoding model. Original optical flow experienced (a) encoded by 26 LPTCs(b), 3 PCs (c), 6 PCs (d), 10 PCs (e), and 26 PCs (f) and decoded back by Gaussian Process models, whose input size is fitting the number of components, trained on 30% of the drone dataset. The 26 LPTCs allow a systematic reconstruction of the original flow with some artefacts in it. Using 26 PCs provide a high accuracy in the reconstruction of the flow. The reduction of the number of PCs results in a simpler model based on the most Principal Components at the cost of some artefacts. More examples are available in Supplementary Materials. With a similar number of components, PCs provide better results than LPTCs.

In Figure 9, we compare the accuracy of the Gaussian Process model processing LPTC signals on both fly and drone datasets. We observe that increasing the proportion of the training set reduces both the AAE and EPE error dramatically. We observe that the Gaussian Model optimises the results globally without favouring the angle or the magnitude of the local flow, as shown with the ratio of EPE and AAE kept mostly constant, no matter the size of the training set. However, we observe that fitting the model on drone data requires a significant proportion of instances in the training set, reducing the EPE to a significant level but still facing difficulties in estimating correctly the local orientation of the vectors. Particularly, the median angular error, as well as its first and third quartile are reduced to a low level but numerous outliers remain present within the resulting maps, as indicated by the high values of the top limit of the box plot located at 1.5x the inter-quartile range of the errors. On the other hand, the decoding of optical flow maps from fly trajectories converges rapidly and reaches a plateau with a training set representing solely 30% of the dataset. Figure 11 highlights the average error within the field of view for drone and fly trajectories. The average angular error for fly trajectories after training of 90% of the dataset is mostly located vertically around ±90 degrees and beyond ±150 degrees azimuth. We found that some outlier vectors located in this region were giving a largely incorrect direction to what is expected thus dramatically affecting the average estimate.

**Figure 9:**
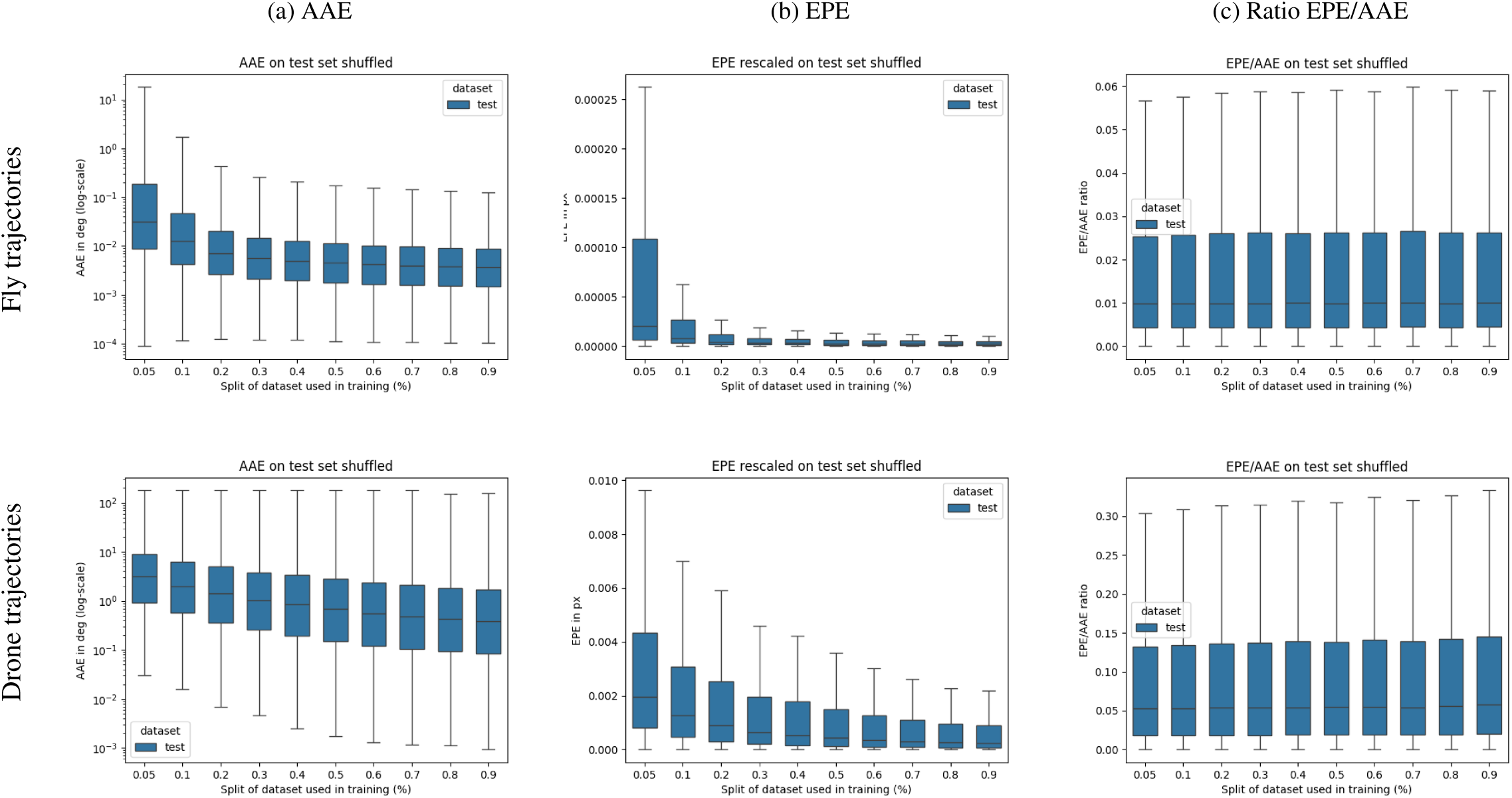
Comparison of accuracy of the Gaussian Process model on fly trajectories (top) and drone trajectories (bottom).

These errors, located in the 25th, 75th, and 99th percentiles of the data, are shown in Figure 10. The 25th percentile map highlights a location of errors a large azimuth, similar to the location present in the average error map. This indicates that this region is prone to frequent but small error. The 75^th^ percentile highlights a second location at zero azimuth and 75 degrees elevation also presents in the average error map. This indicates another frequent location of error, most likely coming from the model itself. Finally, the 99th percentile error map shows outliers randomly distributed over the map.

**Figure 10:**
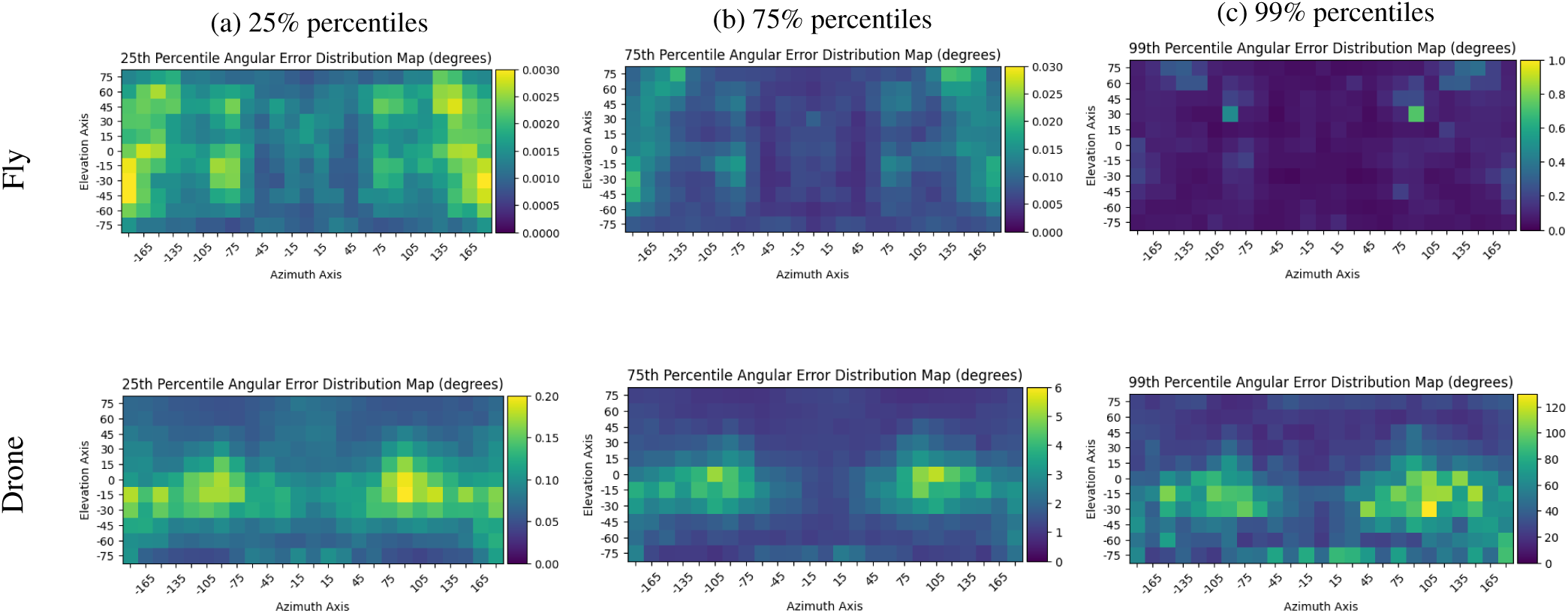
Visualisation of angular error percentiles for fly and drone data. Distribution of the error percentiles (25%, 75%, and 99%) over the field of view for fly (top row) and drone (bottom row) test datasets. Errors have different distributions over the field of view. The distribution of percentiles allow the localisation of errors.

**Figure 11:**
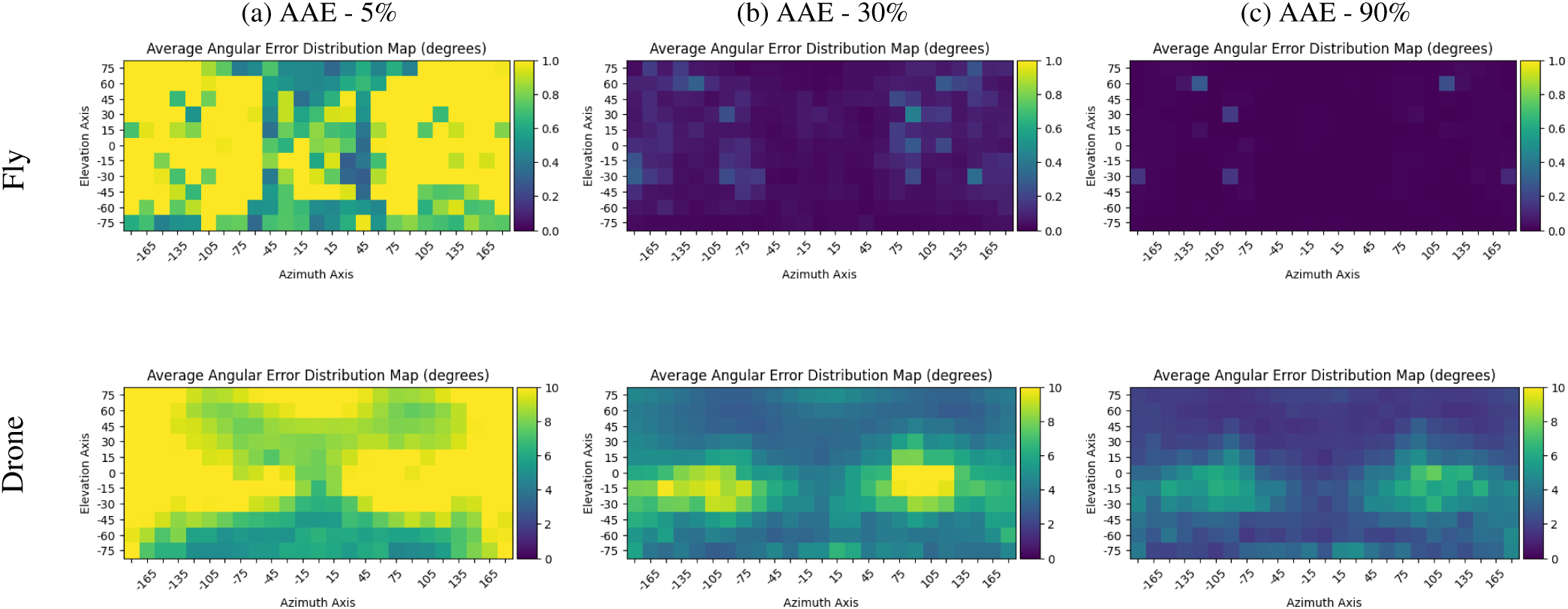
AAE maps based on model evaluation on the test set after being trained on different split of the dataset (5%, 30%, 50%, and 90%). Fly-trained model is show on top row; Drone-based mode is shown on bottom row. The distribution of errors is very different between the two datasets.

Regarding the drone trajectories, we find that the angular errors are mostly distributed horizontally around the ± 90 degrees azimuth axis, describing difficulties to estimate the orientation of the flow in pitch-dominant scenarios, particularly present in the drone trajectories of FlyView which were captured with a quadcopter pitching down when performing a forward translation. Additionally, the EPE is the highest around these specific azimuth points (See Figure 12), as well as at the bottom of the flow where the magnitude is usually the largest due to the forward motion as shown in Figure 1), confirming the difficulty to estimate the flow accurately as a whole and not specifically failing over the magnitude or the vector orientation. As a result, the LPTCs do not fit well the dynamics of the drone and they are not optimally adapted to encode the optical flow generated by its motion, leading to strong errors.

**Figure 12:**
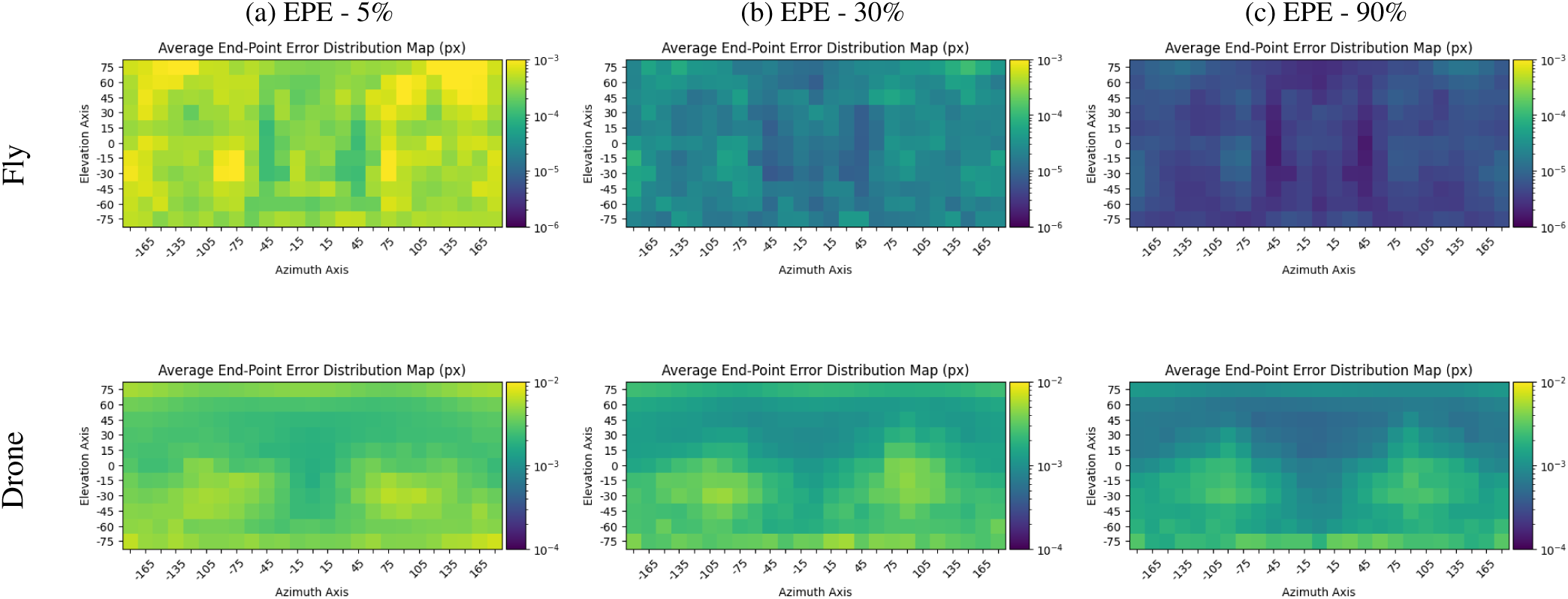
EPE maps based on model evaluation on the test set after being trained on different split of the dataset (5%, 30%, 50%, and 90%). Fly-trained model is show on top row; Drone-based mode is shown on bottom row. Due to the large difference of magnitude of errors, both datasets are not shown with the same range of errors.

Using a Gaussian Process model with Principal Component outputs (see Figure 8), we evaluated the model’s performance with a reduced number of components. When relying on 26 PCs, we observed a slightly more accurate reconstruction of the optical flow compared to the 26 LPTCs, with fewer outliers. However, artefacts were still observed, particularly in regions where the original optical flow exhibited discontinuities in the vector field. These discontinuities, induced by the appearance of nearby objects in the translational optical flow component, are further discussed in Section 4. Such outliers were observed in both LPTC- and PC-based reconstructions.

Additionally, when further reducing the number of PCs, the global structure of the original optical flow remained discernible. While using 10 or even 6 PCs often yielded an estimate that aligned well with the original flow field—albeit with significant outliers—using only 3 PCs consistently failed to produce an accurate reconstruction. More examples are given in the Supplementary Materials.

As a result, with an equivalent number of components, the PCs provided a slightly reduced angular error (See Figure 13). Interestingly, increasing the number of component led to an important increase in the global accuracy of the model, reducing the AAE and EPE errors considerably. In addition, it appears that using Principal Components, distributed differently from LPTCs, which are either positioned along the azimuth axis for VS-cells, and along the elevation axis for HS-cells, led to a concentration of errors mostly along the azimuth axis around -15 degree elevation. This phenomenon could highlight the need of templates distributed across the azimuth axis similarly to VS-cells, instead of orthogonal ones as found in PCs, to provide an homogeneously encoding of the rotational components.

**Figure 13:**
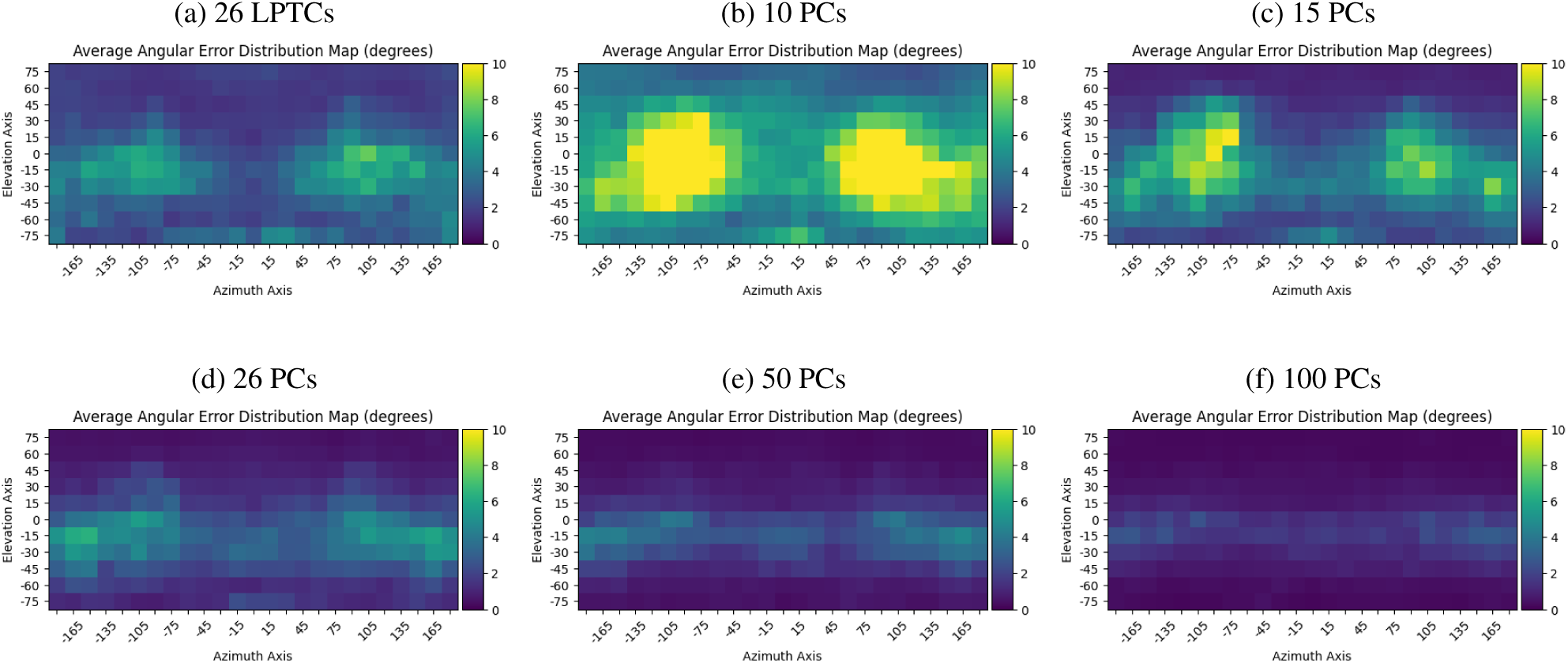
Comparison Angular Error Map between LPTCs-based decoder and PCA-based decoder. Decoder trained on LPTCs (a), or 10(b), 15(c), 26(d), 50(e), and 100(f) PCs with a 90% split of dataset.

## 4 Discussion

While the encoding of optical flow maps describing dynamically-meaningful motions, as found in FlyView, can be effectively modelled using dimensionality reduction techniques, one can wonder about its applicability to a standard optical flow dataset. Specifically, we used the MPI-Sintel dataset [10], a widely used benchmark composed of naturalistic video sequences extracted from open-source animated films. Unlike the natural ego-motions experienced FlyView, the MPI-Sintel dataset features dynamic and non-rigid objects as well as camera movements that simulate cinematographic techniques, such as travelling shots found in long recording sequences.

Similar to the FlyView data, the first principal components derived from the MPI-Sintel dataset exhibited coherent patterns. These components largely corresponded to global motion patterns, such as camera translations which can be observed in the dataset’s cinematographic travelling sequences (see Figures and 14c). Additionally, we observed that only a limited number of principal components were sufficient to explain most of the variance in the dataset. This observation aligns with findings from the FlyView data, where dimensionality reduction techniques reached a practical plateau in capturing variance when increasing input resolution. This limit is similarly reflected in the MPI-Sintel dataset, as shown in Figure 14a.

**Figure 14:**
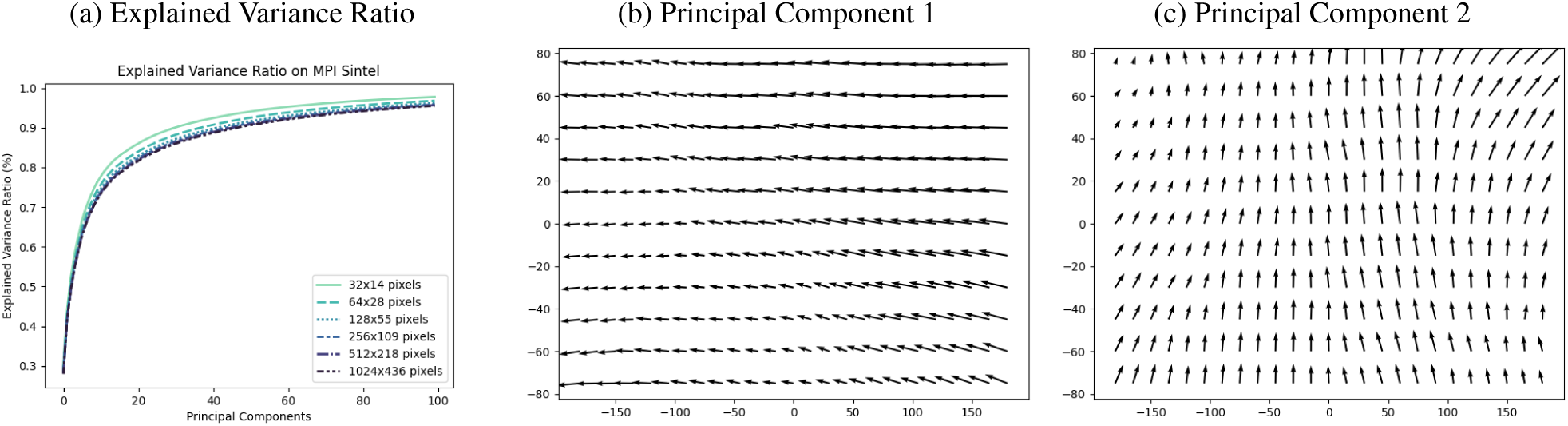
Explained Variance Ratio of PCA components with respect to the resolution of MPI-SIntel and templates from the two principal components.

Despite the similarities, we observe key differences that indicate the limitations of PCA for encoding detailed information in complex visual environments. While global response maps from PCA, akin to the fly’s tangential cells, provide a broad understanding of motion experienced by the fly or the camera, they inherently overlook finer details present in dynamic scenes. For example, the first few principal components in the MPI-Sintel dataset and the 26 LPTCs in the fly focus on capturing global motion but may fail to encode critical information necessary for specific behavioural tasks, such as hunting prey, mating, or avoiding looming obstacles. Figures 21 and 23 in Supplementary Materials illustrate scenarios in which discontinuity was found in the original optical flow maps. After being encoded using either LPTCs or different number of PCs, Gaussian Process models failed not only to accurately reconstruct the area of strong discontinuity, but also introduced numerous artefacts surrounding the discontinuity. Similarly, Figure shows that Principal Components have more difficulties to reach the 100% of explained variance in MPI-SIntel, indicating limitations in capturing the essence of optical flow maps with dynamical objects. Such scene require a more detailed description, especially in dynamic environments featuring objects such as trees, grass, or predators. Nevertheless, species with a limited number of ommatidia, such as *Drosophila* are unlikely to be able to detect small and dynamic objects in their field of view as the low number of photoreceptors is providing a blurred and low resolution view of the scene.

To address these limitations in species whose number of ommatidia is higher, it is relevant to consider the role of complementary motion-sensitive inputs, such as cells tuned to looming stimuli. Studies have shown that loom-sensitive neurons in the fly, such as those described by [65] and [39], play an essential role in detecting imminent collisions or rapidly approaching objects. These cells might work in collaboration with VS and HS cells to refine the fly’s understanding of its surroundings. Indeed, while VS and HS cells encode global optical flow patterns for navigation, loom-sensitive neurons add an additional layer of spatial and temporal resolution, allowing the fly to react to immediate threats or opportunities. This division of labour among different motion-sensitive neural pathways enables the fly to efficiently process complex visual environments and respond appropriately to both global and localized stimuli. However, the study of the two categories of cells could show that they complement one-another thus permitting loom sensitive cells to refine the pose estimation task.

Analogously to the increase of performance with respect to the number of PCs, biological systems might benefit from an increased number of motion-sensitive neurons with different template. This could explain why *Drosophila*, *Musca domestica*, and *Calliphora* exhibit different numbers of VS-cells, potentially allowing species to use more complex flight dynamics thanks to their improved encoding capabilities processing more complex optical flow.

Discontinuities in the optical flow field, particularly those induced by environmental discontinuities such as object boundaries or sudden motion changes, introduce artefacts that could affect the stability of motion perception. H- and V-cells serve as prime candidates in mitigating these distortions, acting as a form of signal correction mechanism that ensures the consistency of the motion signal, irrespective of environmental obstacles, in both hemispheres. By stabilizing motion signals, these cells could contribute to more robust optical flow processing, reducing the impact of dynamic environmental changes and observed obstacles.

Additionally, other motion-sensitive neurons, such as the CH- and FD-cells, may complement this process. CH neurons have been identified as wide-field inhibitors responsible for the small-field tuning of the figure-detection neuron FD1 [15, 19]. This interaction plays a critical role in figure-ground discrimination, a fundamental task in visual motion processing. The ability of FD-cells to compensate for the limitations of HS-cells in distinguishing between object and background motion further suggests a specialized division of labour among motion-sensitive neurons [15–17].

Taken together, we hypothesize that the integration of different motion-sensitive neurons, including VS, HS, V, and H cells, as well as CH cells and figure-detection neurons FD-cells, may form a multi-layered processing system that allows the fly to construct an internal obstacle-free image of the optical flow in parallel to obstacle detection, solely used as a filtered signal for self-motion.

Extending this idea, the integration of complementary motion-processing strategies could enhance computational models of optical flow encoding, particularly for tasks requiring fine-grained scene understanding. For example, incorporating features inspired by looming-like optical flow into the optical flow templates used for encoding might improve their ability to capture both global patterns and localized motion cues. Additionally, Tanaka et al. [67] identified some components of *Drosophila*’s circuitry for stationary pattern detection, which run in parallel to the LPTCs. Considering the limitations of global optical flow templates, such a parallel network might also be of inspiration to fully capture the environment of the observer. Such advancements would be particularly beneficial for applications involving dynamic environments, such as autonomous navigation, where balancing global motion encoding with local detail detection is critical.

The LPTCs resulted in accurate representation of the experienced flow on the quadcopter trajectories from FlyView [47]. However, the specialization of LPTCs for natural fly motion highlights their evolutionary advantage in extracting biologically relevant visual cues. LPTCs embed a strong prior on the optical flow distributions typical of naturalistic fly trajectories, enabling them to process a well-defined range of motion patterns efficiently. When a quadcopter undergoes motion dynamics outside this range—such as abrupt translations, rotations, or non-biological manoeuvres—the ability to decode motion or reconstruct the original optical flow becomes less reliable. This limitation suggests that the LPTC system is optimized for the fly’s navigation demands rather than for general motion encoding across diverse dynamics.

Interestingly, the description of coherent patterns close to the ones generated by pure rotations or translations suggests that not only Principal Components are a prime candidate for pose estimation tasks in robotics, but also that a selected number of PCs could be enough to provide a full 6-DoF pose estimate. As a matter of fact, the first three PCs, PC1, PC2, PC3, describe the main rotation axes, Yaw, Pitch, and Roll respectively. These patterns, found when processing the full field of view with fly data, but also when splitting the hemispheres in fly and drone data, indicate that rotations are, for agents of different dynamics, the motions where most of the energy is capture. Additionally, both PC1 and PC2 include a small component of Roll rotation, which is mostly encoded by PC3. Whilst a separate response of PC1 and PC2 would provide information on Yaw and Pitch rotations, a compound response of both PCs could provide some Roll information. Nevertheless, even if such a rotation can be estimated without an artificial cell responding specifically to it, having such an additional cell would contribute to remove ambiguities in some scenarios Particularly, when introducing new motions in the dynamics (e.g. forward displacement) for which a compound response of both PC1 and PC2 is unlikely to be enough to discriminate. However, should the accuracy of a system be the priority, the uncertainty behind a limited number of PCs make the overall system unreliable for critical missions. Additionally, using global optical flow templates, the identification of a 45 degrees azimuth rotation would be more accurately represented by a template specific to this motion rather than an average response of components for Pitch and Roll motions. This is likely explaining why VS cells are densely represented in the Lobula Plate in a non-orthogonal fashion, permitting to identify specific axes of rotation.

Additionally, since Principal Components are orthogonal by definition, a given component should not encode a combination of distinct motions unless those motions are inherently associated. Notably, we observed in the Results section that the yaw-dominated PC1 contains a measurable roll component, while the roll-dominated PC3 also includes a degree of yaw. These combined rotational elements are likely reflective of the manoeuvres performed by the blowfly during turns, which may serve to minimise the energetic cost of correcting its heading. Similar coupling has been observed in vision-based flight control in the hawkmoth *Hyles lineata* [74], suggesting that the blowfly may likewise produce coupled roll and yaw moments in response to yaw stimuli.

In robotic systems, optical flow is frequently used for maintaining altitude, stabilizing flight, and detecting obstacles. However, unlike biological systems, which naturally encode motion patterns through specialized neurons, artificial systems often rely on computationally expensive algorithms to extract optical flow features and estimate six degrees of freedom (6-DoF) pose. Incorporating bio-inspired solutions such as LPTCs could bridge this gap by enabling lightweight and efficient motion encoding. As a result, artificial LPTCs (a-LPTCs) could be designed to specialize in the motion dynamics of specific platforms, such as quadcopters. These a-LPTCs would embed priors tailored to the quadcopter’s operational motion range and flight characteristics, allowing for more efficient encoding of visual information. By mimicking the distributed processing architecture of biological LPTCs, artificial systems could enhance performance in tasks such as collision avoidance, path planning, and target tracking—especially in environments with complex optical flow patterns. Additionally, combining these artificial structures with advanced machine learning techniques—such as artificial neural networks—could further enhance their adaptability to diverse environments and motion dynamics, opening the way to low-energy self-motion estimation in real time and in complex environments.

Unlike Principal Components, which are limited to orthogonal axes and provide only three independent rotational components for linear encoding, LPTCs encode partial rotations in a higher number, allowing for non-linear motion representation. Additionally, beyond PC8, the extracted Principal Components exhibit incoherent patterns, limiting their usefulness for accurate motion reconstruction and interpretability. The best representation of a-LPTCs may therefore involve using a larger number of coherent templates, similar to LPTCs, but with the encoding advantages of PCs—such as spanning the entire field of view (FoV) with both positive and negative responses. This would enable finer reconstruction of optical flow while maintaining computational efficiency.

A first step in designing such a system could involve using the Preferred Direction flow maps [48] of biological LPTCs as matched filters to extract relevant motion patterns. A second step would involve designing a set of templates whose rotation axes are spatially distributed across the FoV. These templates could follow either a uniform distribution for general scenarios or be fine-tuned to the specific motion dynamics of the platform, optimizing state estimation and optical flow reconstruction.

Finally, the Gaussian Process model was demonstrated to be a first success in decoding optical flow from vector field templates. A similar methodology could potentially be applied to environment decoding, analogous to previous hypotheses suggesting that flies extract information about the spatial layout of the environment [37]. Therefore, future studies could explore this avenue to refine our understanding of the fly builds an internal representation of its surrounding environment based on structure-from-motion estimated from encoded optical flow information.

## 5 Conclusion

In our study, we explored the principles of dimensionality reduction underlying the functioning of Lobula Plate Tangential Cells. Through a series of experiments using data-driven dimensionality reduction and reconstruction techniques applied to an extensive database of trajectories from flies and drones, we extracted critical insights from the encoding of visual motion by LPTCs.

To the best of our knowledge, this is the first time a big data approach has been employed to investigate the intricacies of LPTCs response. Our work provides the following contributions to the field. First, we demonstrated, with the example of PCA, that methods based on match global optical flow patterns, serve as an effective means of encoding motion from the environment. These methods yield high explained variance with a minimal number of components. Notably, the primary principal components fitted to natural optical flow experienced during flight correspond to translations and rotations. These findings align with the six degrees of freedom in self-motion, indicating their potential utility in bio-informed engineering applications for navigational tasks.

Secondly, we showed that LPTCs, despite being a destructive dimensionality reduction mechanism, are capable of compressing optical flow information which can then be reconstructed thanks to priors and the complementarity of hemispheres. By employing Gaussian Processors for a first reconstruction, we opened the door to more efficient and accurate methods, such as deep neural networks, which could further enhance the fidelity of optical flow reconstruction.

Finally, we highlighted that while global optical flow patterns provide a robust framework for encoding optical flow for navigation tasks, they are unlikely to be effective for applications requiring precise local vector accuracy. This limitation becomes apparent in scenarios involving non-continuous displacements within the optical flow, such as those generated by dynamic objects or prey movements.

## Acknowledgments and Disclosure of Funding

AL is supported by a doctoral studentship funded by Dstl, the Department of Biology, and Jesus College, Oxford. This project has received funding from the European Research Council (ERC) under the European Union’s Horizon 2020 research and innovation programme (grant agreement No. 682501). We thank Holger G Krapp for his useful recommendations and insightful discussions on fly vision as well as for sharing the LPTC recordings of the 13 LPTC Calliphora cells of the right hemisphere. We declare no competing interests. This document is an overview of UK MOD’s Defence Science and Technology Laboratory (Dstl) sponsored research and is released for informational purposes only. The contents of this document should not be interpreted as representing the views of the UK MOD, nor should it be assumed that they reflect any current or future UK MOD policy.

## Supplementary Materials

### Explained Variance Ratio of Principal Components

**Table 1:** Explained Variance for Principal Components on Right hemisphere with mirrored data.

|  | **25x11px** | **50x22px** | **100x45px** | **250x110px** |
| --- | --- | --- | --- | --- |
| **PC1** | 0.435624 | 0.426979 | 0.407201 | 0.391816 |
| **PC2** | 0.726102 | 0.715985 | 0.689284 | 0.66867 |
| **PC3** | 0.833775 | 0.810255 | 0.786473 | 0.767282 |
| **PC4** | 0.90500 | 0.874551 | 0.855426 | 0.834843 |
| **PC5** | 0.949957 | 0.931534 | 0.906858 | 0.883319 |
| **PC6** | 0.977527 | 0.953861 | 0.927937 | 0.908244 |
| **PC7** | 0.984435 | 0.97425 | 0.947805 | 0.928674 |
| **PC8** | 0.989891 | 0.981301 | 0.965661 | 0.946865 |
| **PC9** | 0.992386 | 0.986879 | 0.976016 | 0.958805 |
| **PC10** | 0.993699 | 0.990629 | 0.981013 | 0.967262 |
| **PC11** | 0.994741 | 0.992839 | 0.985766 | 0.971953 |
| **PC12** | 0.995588 | 0.994279 | 0.988205 | 0.976291 |
| **PC13** | 0.9962 | 0.995026 | 0.990287 | 0.979792 |
| **PC14** | 0.996777 | 0.995563 | 0.992128 | 0.98284 |
| **PC15** | 0.997207 | 0.996067 | 0.993427 | 0.98486 |
| **PC16** | 0.997613 | 0.996526 | 0.994202 | 0.98671 |
| **PC17** | 0.997904 | 0.99693 | 0.994906 | 0.988281 |
| **PC18** | 0.99812 | 0.997295 | 0.995368 | 0.989582 |
| **PC19** | 0.998314 | 0.997559 | 0.9958 | 0.990602 |
| **PC20** | 0.998491 | 0.997764 | 0.996206 | 0.991412 |
| **PC21** | 0.998618 | 0.997943 | 0.99658 | 0.992182 |
| **PC22** | 0.998724 | 0.998098 | 0.996906 | 0.992936 |
| **PC23** | 0.998825 | 0.998239 | 0.997166 | 0.993449 |
| **PC24** | 0.998916 | 0.998371 | 0.997419 | 0.993913 |
| **PC25** | 0.998997 | 0.998479 | 0.997603 | 0.994344 |
| **PC26** | 0.999069 | 0.998574 | 0.997763 | 0.994747 |

**Table 2:**
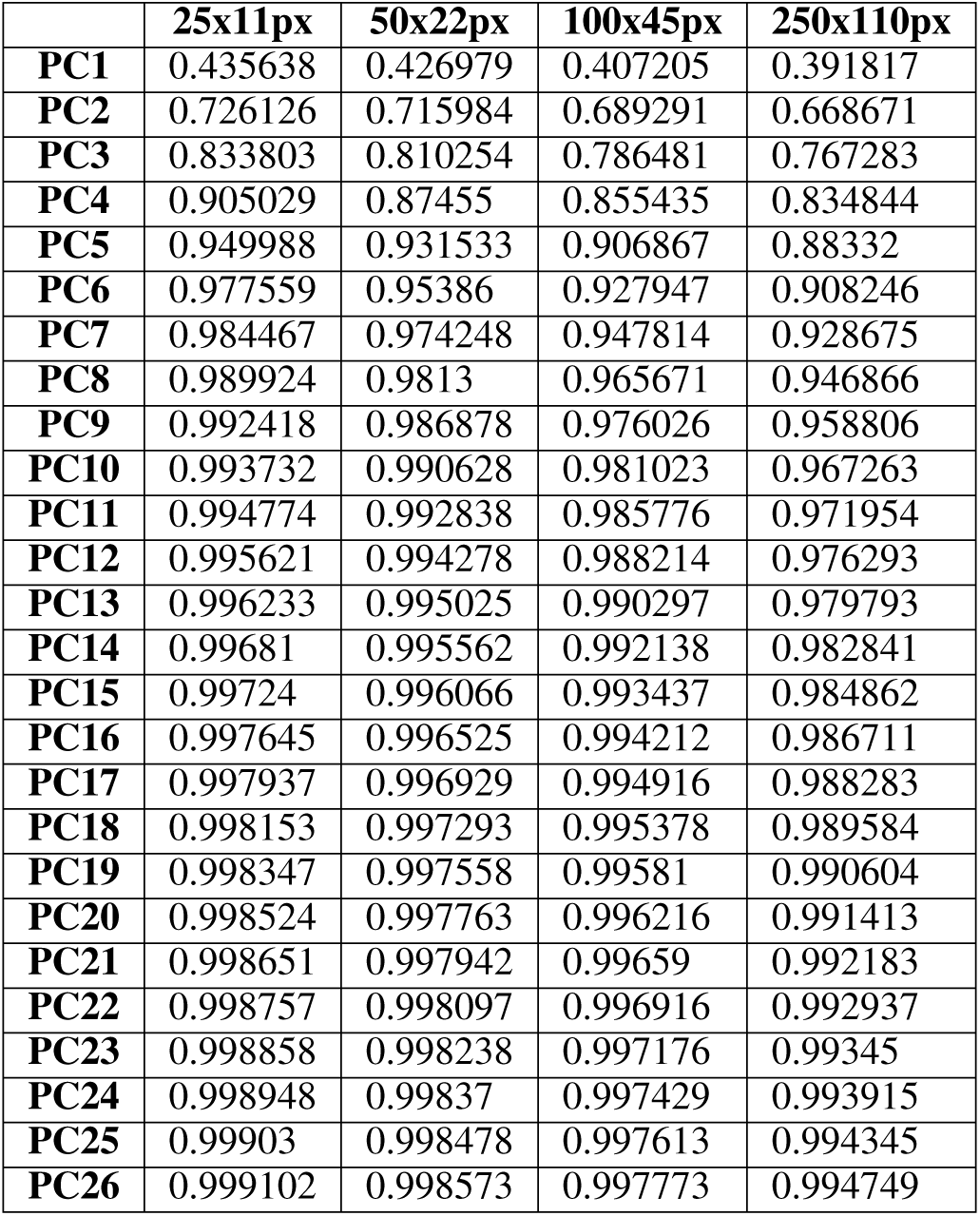
Explained Variance for Principal Components on Left hemisphere with mirrored data.

|  | **25x11px** | **50x22px** | **100x45px** | **250x110px** |
| --- | --- | --- | --- | --- |
| **PC1** | 0.435638 | 0.426979 | 0.407205 | 0.391817 |
| **PC2** | 0.726126 | 0.715984 | 0.689291 | 0.668671 |
| **PC3** | 0.833803 | 0.810254 | 0.786481 | 0.767283 |
| **PC4** | 0.905029 | 0.87455 | 0.855435 | 0.834844 |
| **PC5** | 0.949988 | 0.931533 | 0.906867 | 0.88332 |
| **PC6** | 0.977559 | 0.95386 | 0.927947 | 0.908246 |
| **PC7** | 0.984467 | 0.974248 | 0.947814 | 0.928675 |
| **PC8** | 0.989924 | 0.9813 | 0.965671 | 0.946866 |
| **PC9** | 0.992418 | 0.986878 | 0.976026 | 0.958806 |
| **PC10** | 0.993732 | 0.990628 | 0.981023 | 0.967263 |
| **PC11** | 0.994774 | 0.992838 | 0.985776 | 0.971954 |
| **PC12** | 0.995621 | 0.994278 | 0.988214 | 0.976293 |
| **PC13** | 0.996233 | 0.995025 | 0.990297 | 0.979793 |
| **PC14** | 0.99681 | 0.995562 | 0.992138 | 0.982841 |
| **PC15** | 0.99724 | 0.996066 | 0.993437 | 0.984862 |
| **PC16** | 0.997645 | 0.996525 | 0.994212 | 0.986711 |
| **PC17** | 0.997937 | 0.996929 | 0.994916 | 0.988283 |
| **PC18** | 0.998153 | 0.997293 | 0.995378 | 0.989584 |
| **PC19** | 0.998347 | 0.997558 | 0.99581 | 0.990604 |
| **PC20** | 0.998524 | 0.997763 | 0.996216 | 0.991413 |
| **PC21** | 0.998651 | 0.997942 | 0.99659 | 0.992183 |
| **PC22** | 0.998757 | 0.998097 | 0.996916 | 0.992937 |
| **PC23** | 0.998858 | 0.998238 | 0.997176 | 0.99345 |
| **PC24** | 0.998948 | 0.99837 | 0.997429 | 0.993915 |
| **PC25** | 0.99903 | 0.998478 | 0.997613 | 0.994345 |
| **PC26** | 0.999102 | 0.998573 | 0.997773 | 0.994749 |

**Table 3:** Explained Variance for Principal Components on both hemispheres combined with mirrored data.

|  | **25x11px** | **50x22px** | **100x45px** | **250x110px** |
| --- | --- | --- | --- | --- |
| **PC1** | 0.394067 | 0.37021 | 0.351837 | 0.339715 |
| **PC2** | 0.708929 | 0.687174 | 0.659699 | 0.639533 |
| **PC3** | 0.798912 | 0.77313 | 0.747272 | 0.726462 |
| **PC4** | 0.86146 | 0.825378 | 0.807306 | 0.788186 |
| **PC5** | 0.900586 | 0.874631 | 0.85137 | 0.829693 |
| **PC6** | 0.937252 | 0.916585 | 0.89131 | 0.867803 |
| **PC7** | 0.959692 | 0.941979 | 0.914314 | 0.889493 |
| **PC8** | 0.972766 | 0.957826 | 0.931501 | 0.908204 |
| **PC9** | 0.985598 | 0.967893 | 0.944011 | 0.923971 |
| **PC10** | 0.98858 | 0.975821 | 0.955197 | 0.936644 |
| **PC11** | 0.991216 | 0.983179 | 0.964225 | 0.946197 |
| **PC12** | 0.992639 | 0.986208 | 0.972659 | 0.954435 |
| **PC13** | 0.993639 | 0.989003 | 0.979509 | 0.962115 |
| **PC14** | 0.994486 | 0.991027 | 0.982285 | 0.966867 |
| **PC15** | 0.995172 | 0.993018 | 0.984969 | 0.971317 |
| **PC16** | 0.995703 | 0.993749 | 0.986809 | 0.97434 |
| **PC17** | 0.996208 | 0.99442 | 0.988569 | 0.977265 |
| **PC18** | 0.996666 | 0.994958 | 0.990125 | 0.979363 |
| **PC19** | 0.996969 | 0.995471 | 0.991609 | 0.981208 |
| **PC20** | 0.997262 | 0.995893 | 0.992446 | 0.982657 |
| **PC21** | 0.997502 | 0.996296 | 0.993261 | 0.984026 |
| **PC22** | 0.997702 | 0.996678 | 0.99376 | 0.985392 |
| **PC23** | 0.997893 | 0.996926 | 0.994231 | 0.986683 |
| **PC24** | 0.998051 | 0.997141 | 0.994645 | 0.987762 |
| **PC25** | 0.998196 | 0.997341 | 0.995037 | 0.988802 |
| **PC26** | 0.998329 | 0.997509 | 0.995408 | 0.989469 |

### Principal Components on fly trajectories - full field of view

**Figure 15:**
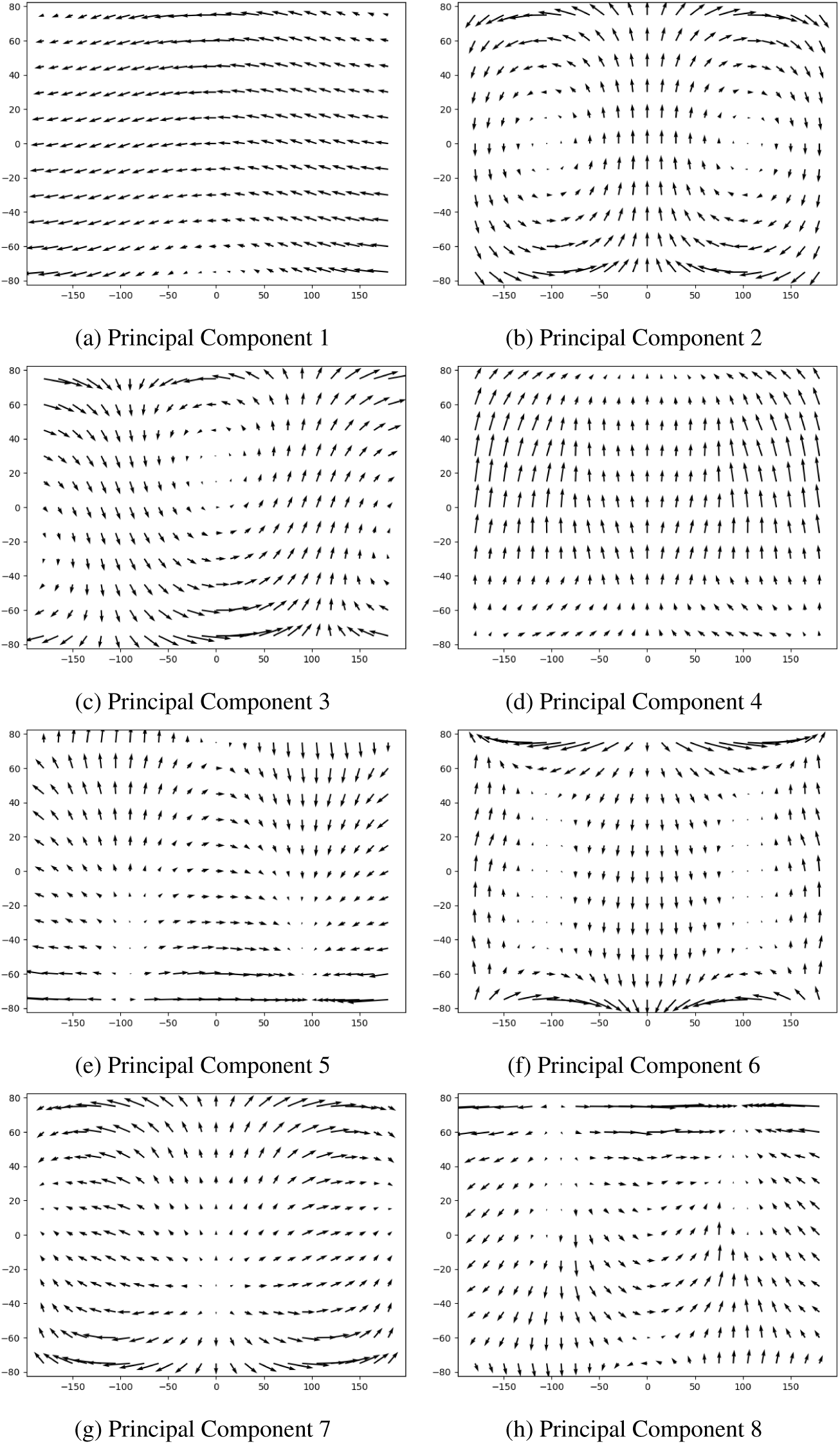
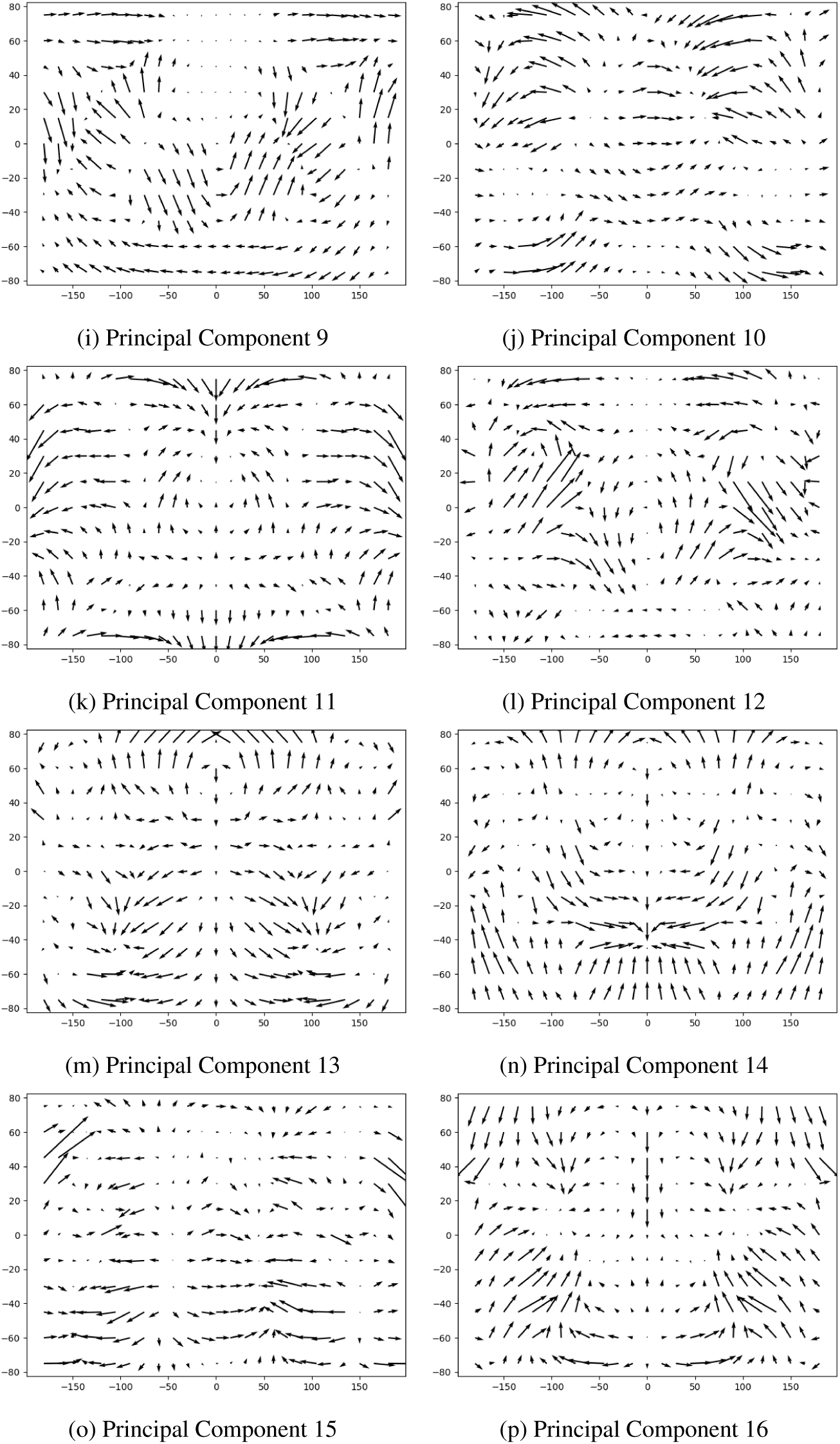
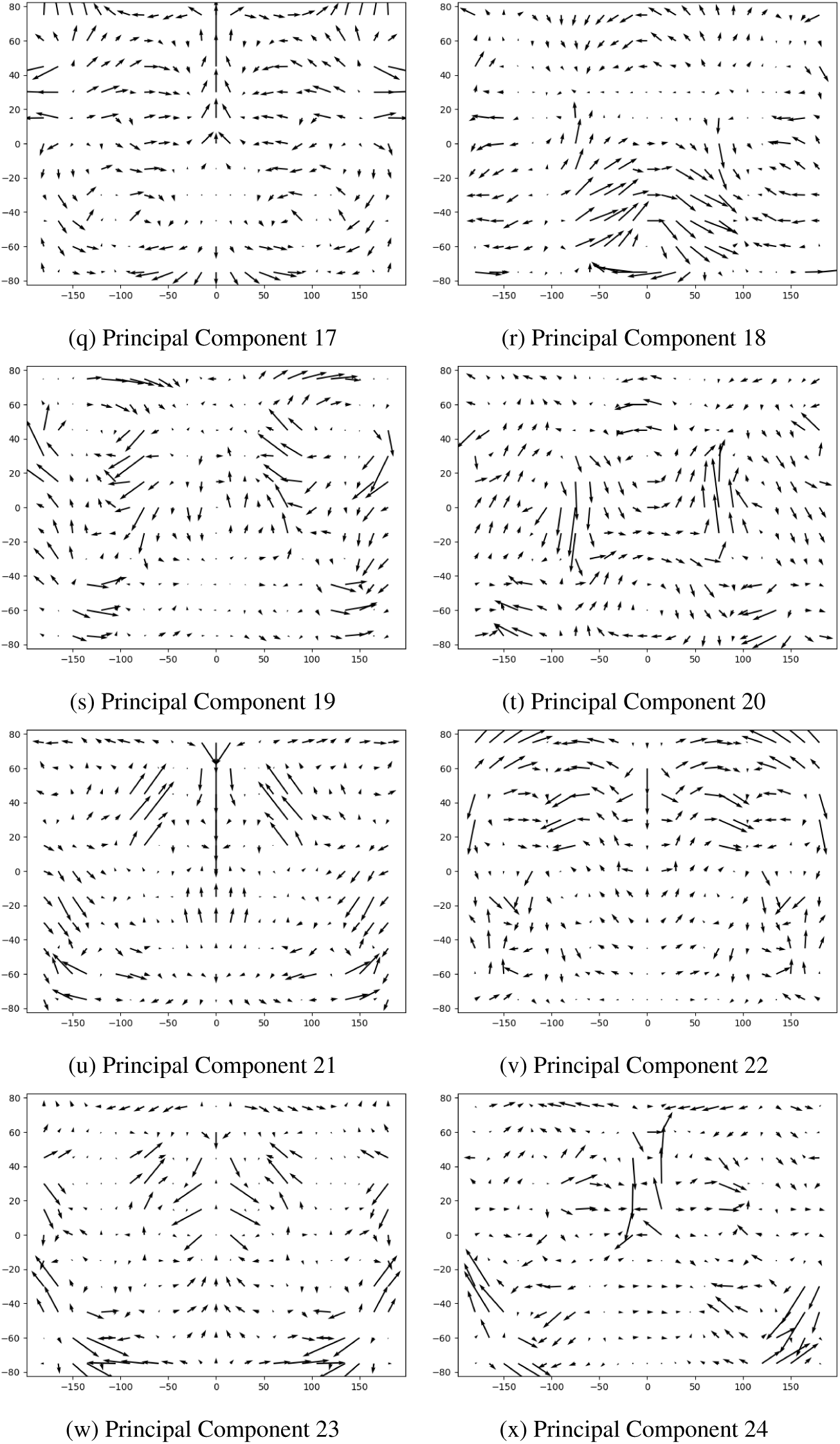
24 First Principal Components fitted to optical flow of fly trajectories from FlyView

### 5.1 Principal Components on fly trajectories - left and right field of view

**Figure 16:**
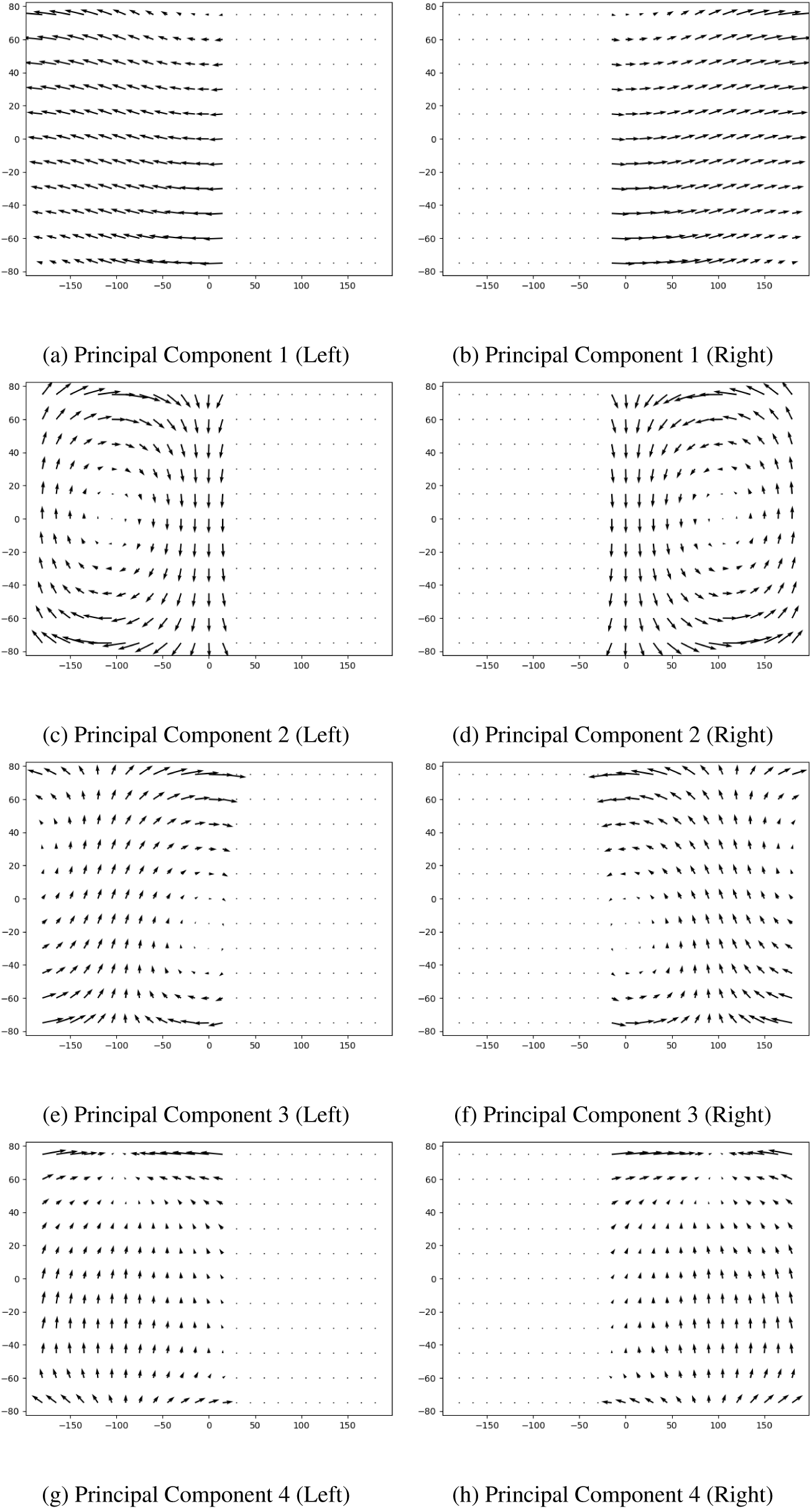
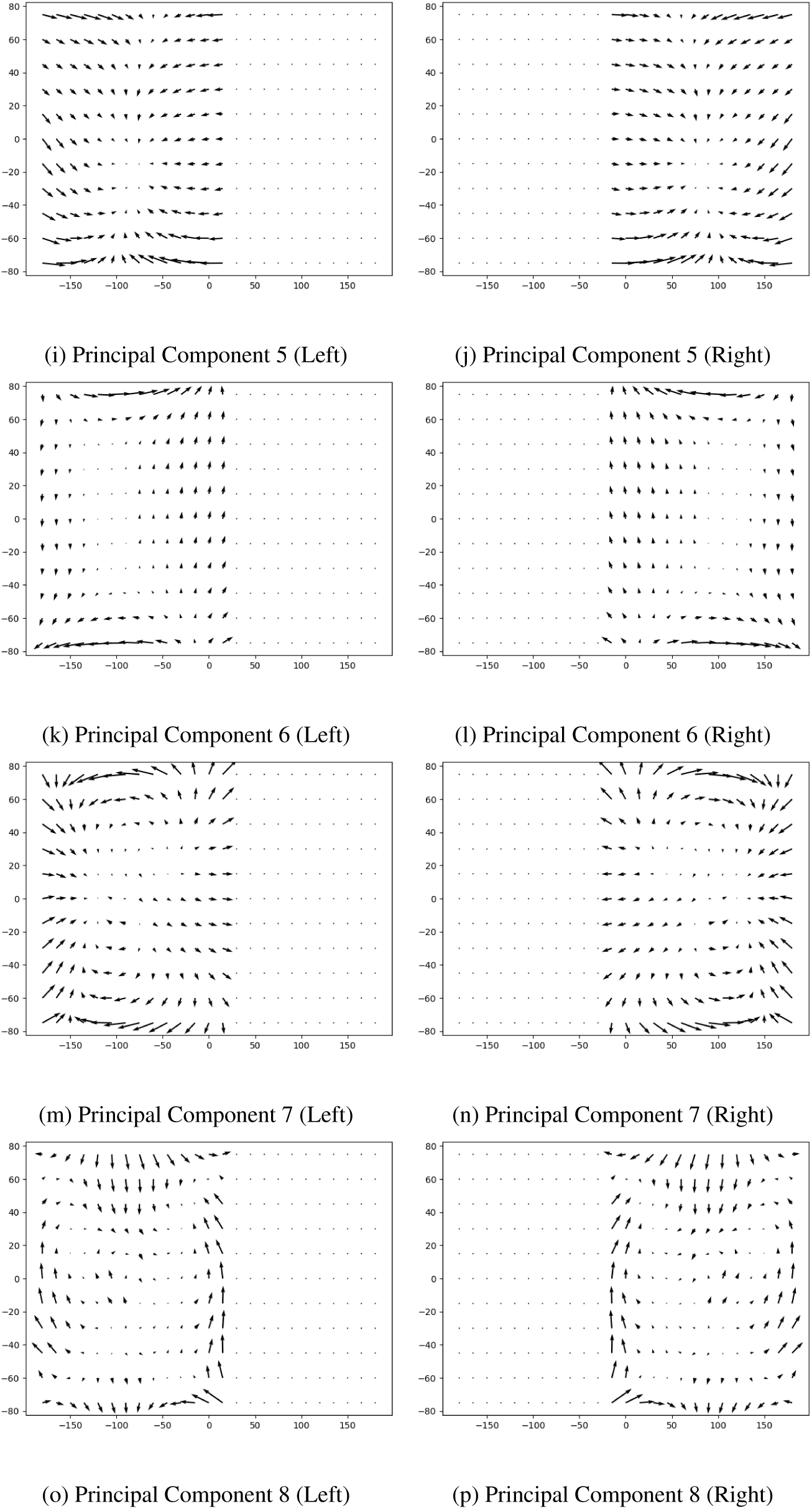
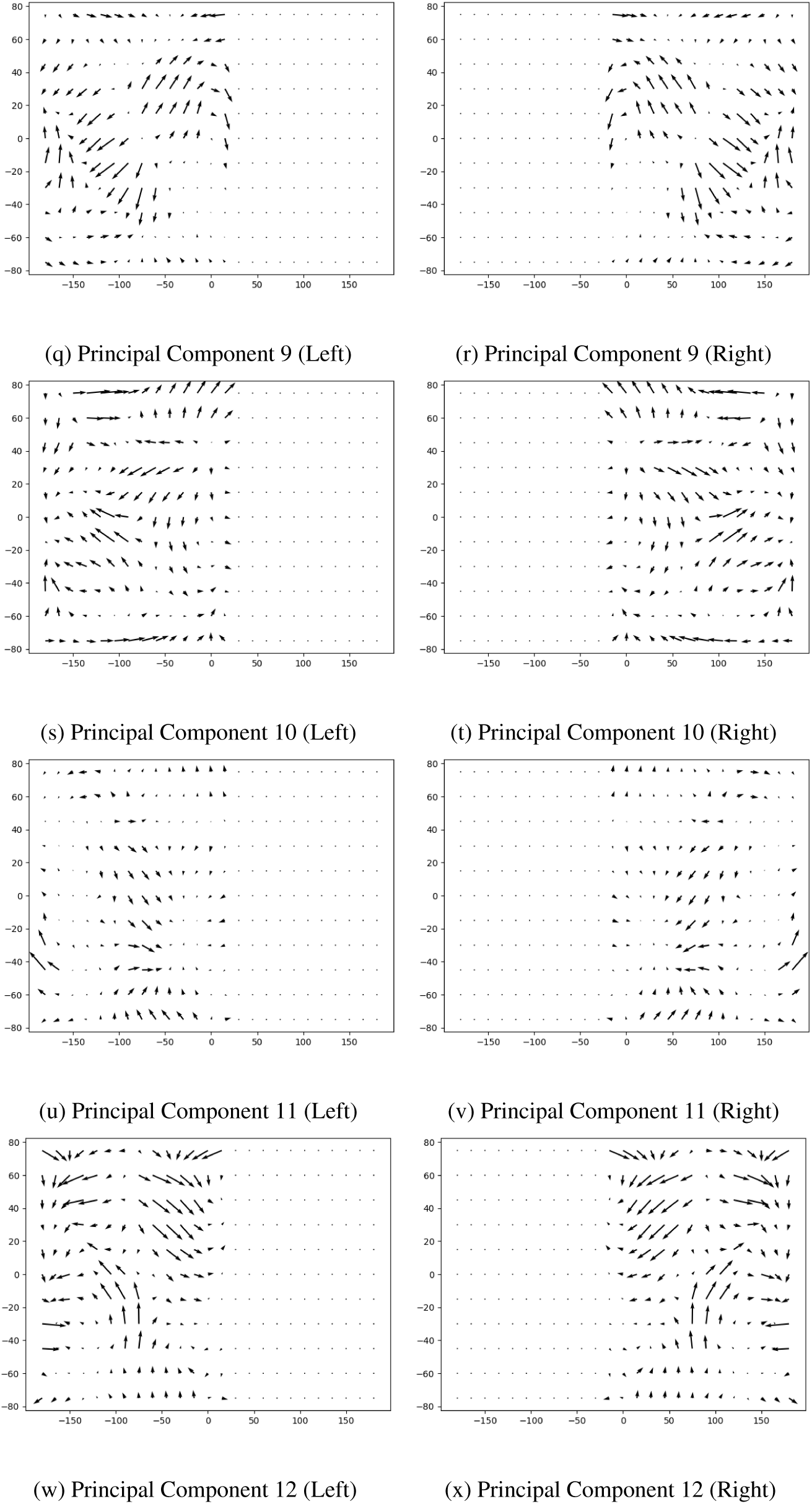
Principal Components on fly trajectories for separated left and right hemispheres

### Principal Components on drone trajectories - left and right field of view

**Figure 17:**
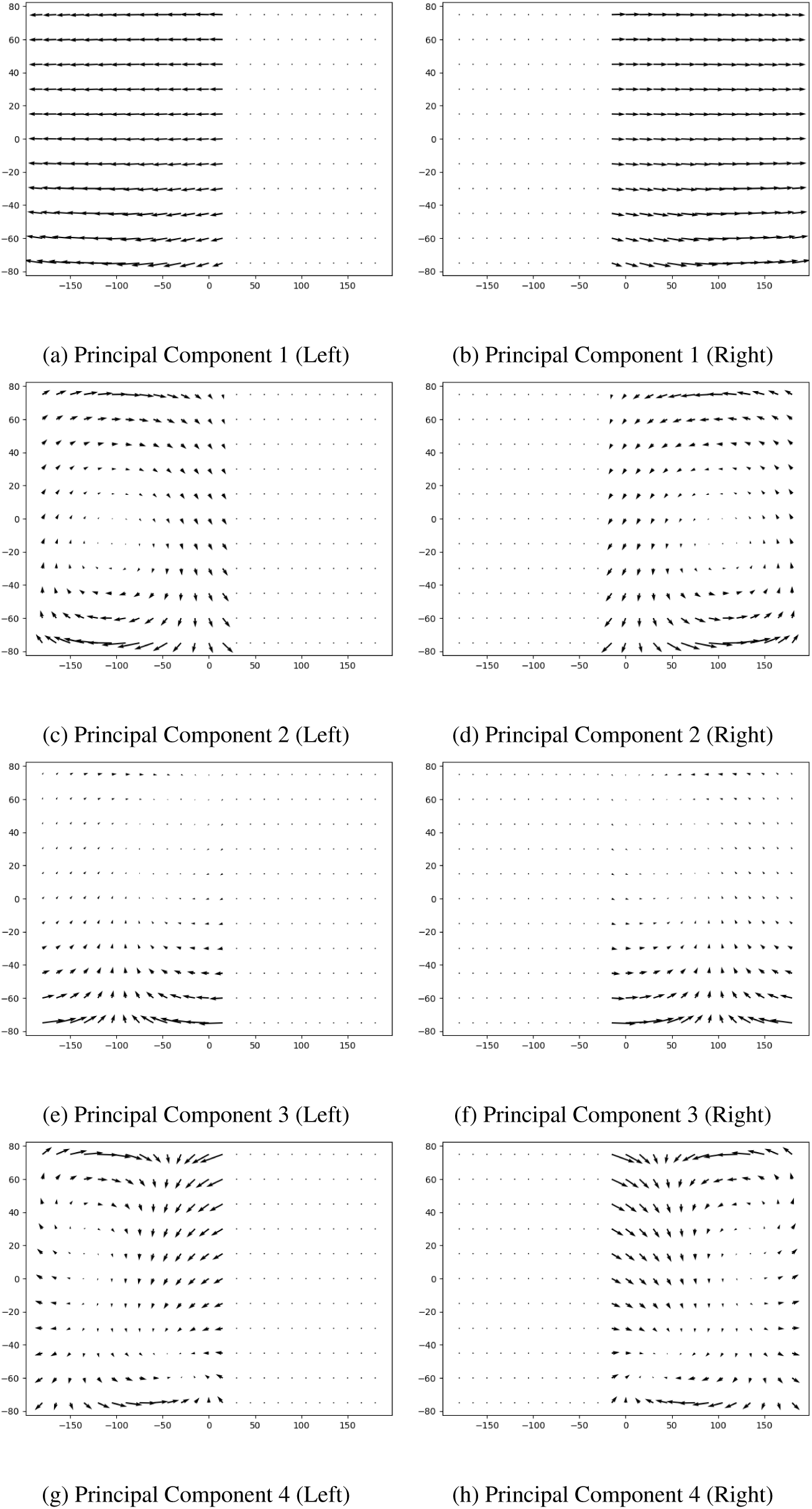
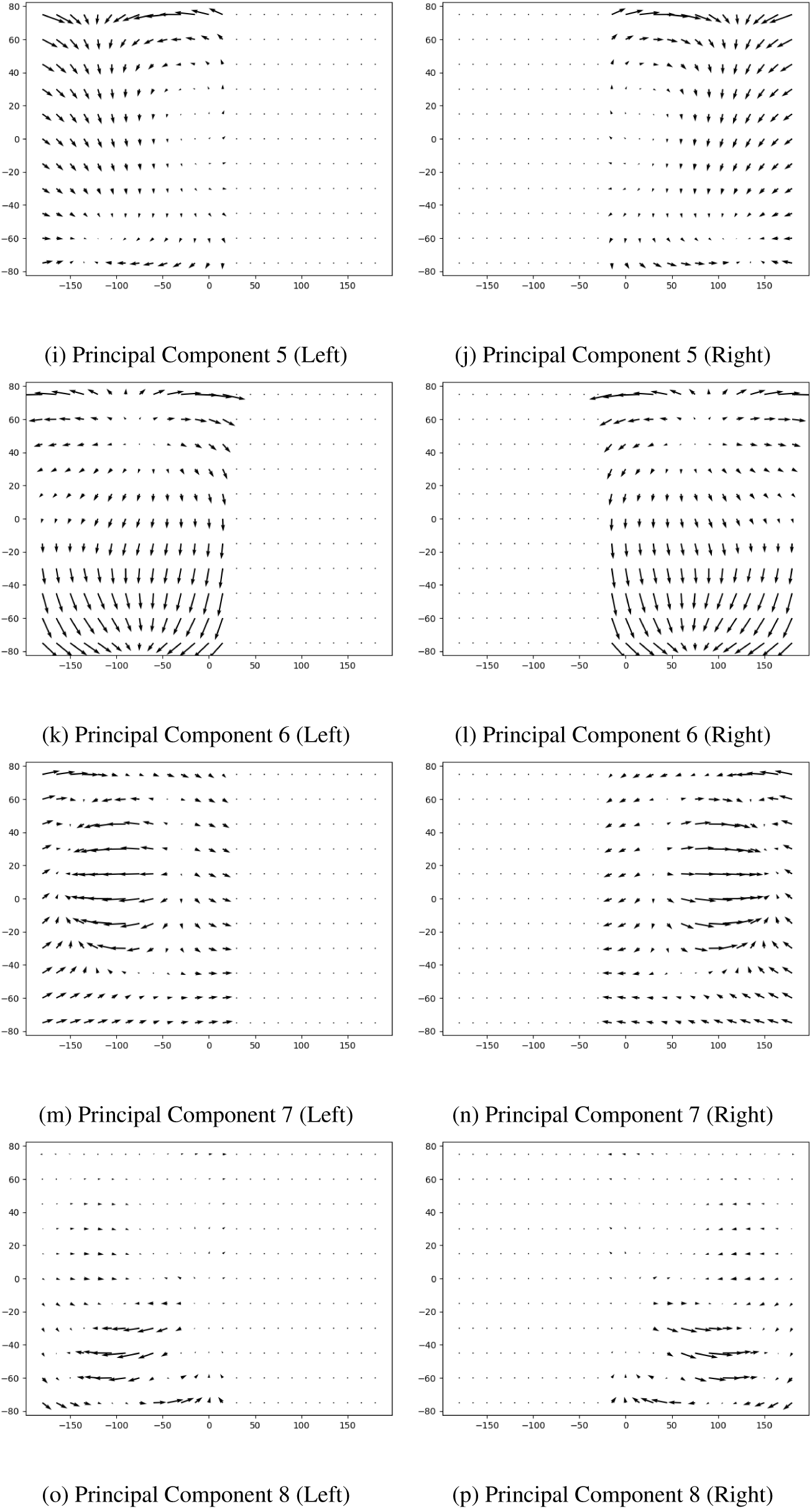
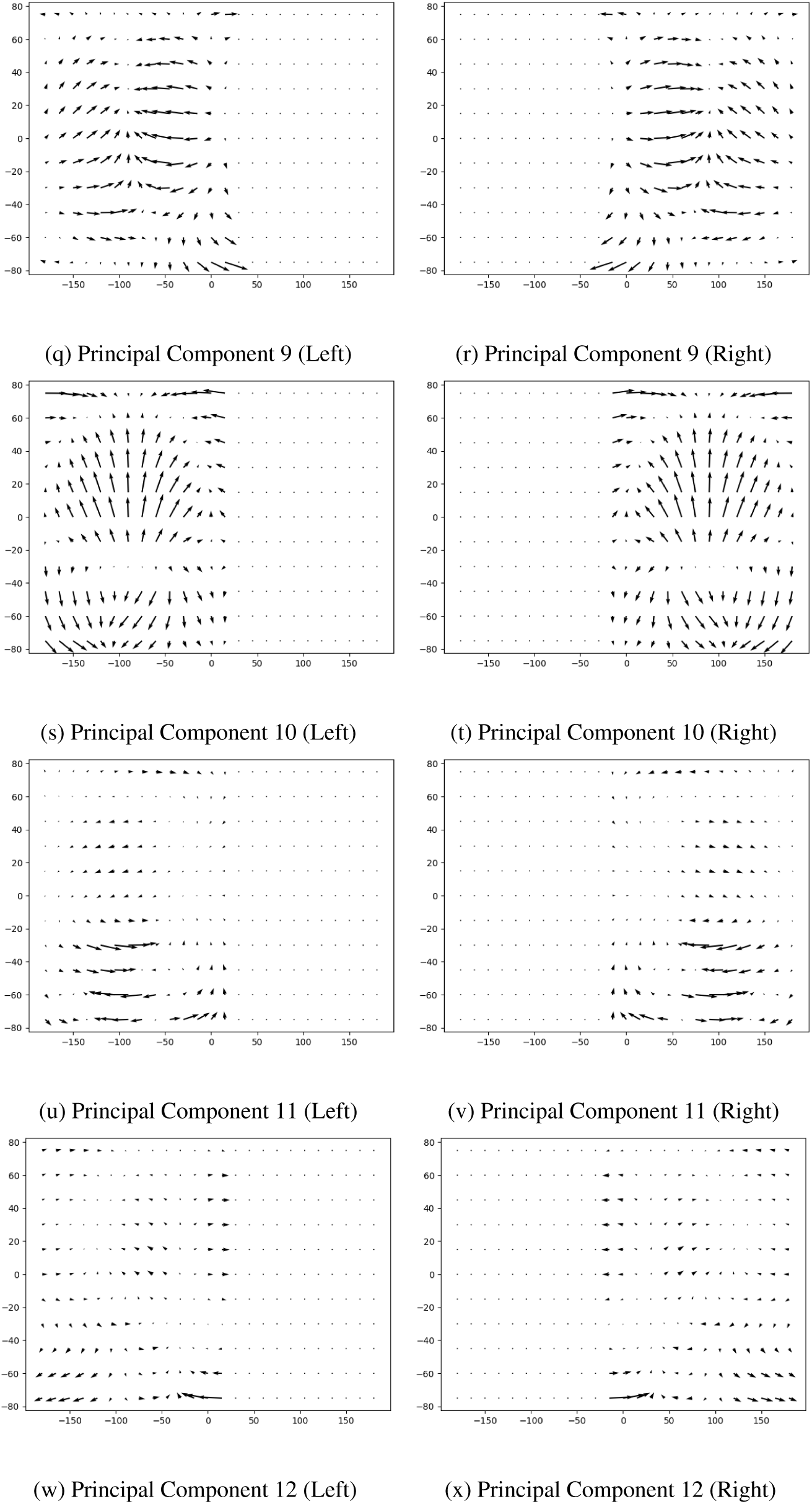
Principal Components on drone trajectories for separated left and right hemispheres

**Figure 18:**
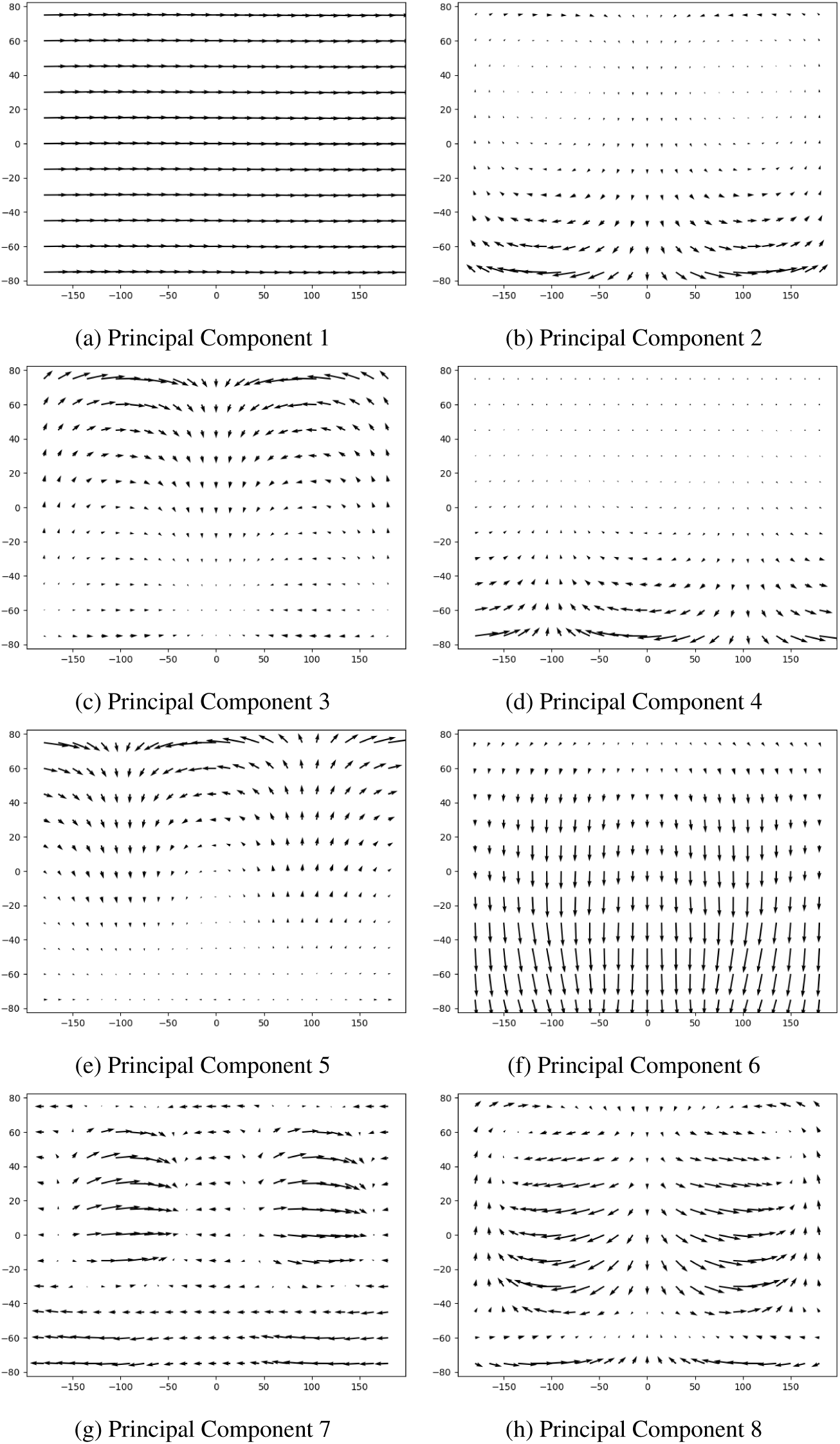
First Principal Components fitted to optical flow of drone trajectories . Drone data from FlyView [47].

**Figure 19:**
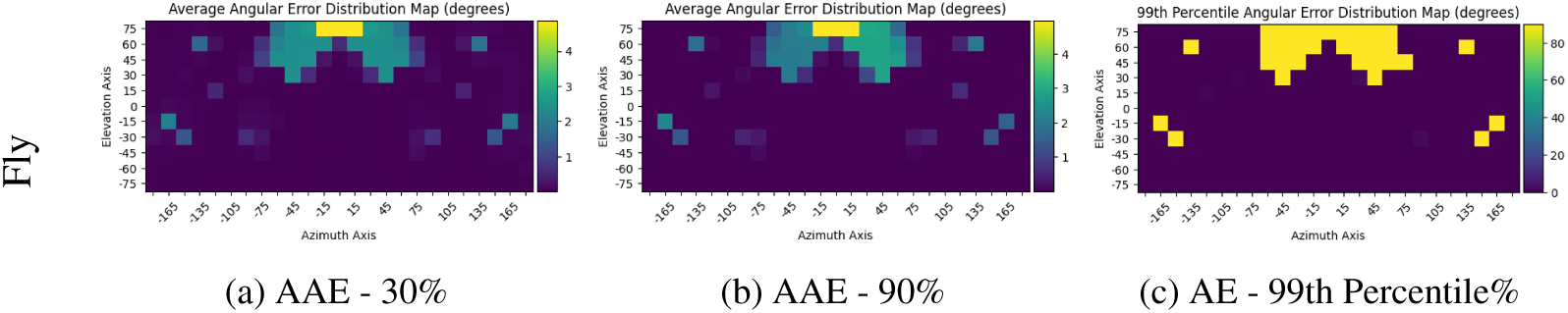
Artefacts in the test set. AAE maps based on model evaluation on the test set after being trained on different split of the dataset (30%, and 90%). Some optical flow maps with zero-valued vectors introduced artefacts in the training leading to outliers with 90 degrees of errors on consistent location in the field of view.

**Figure 20:**
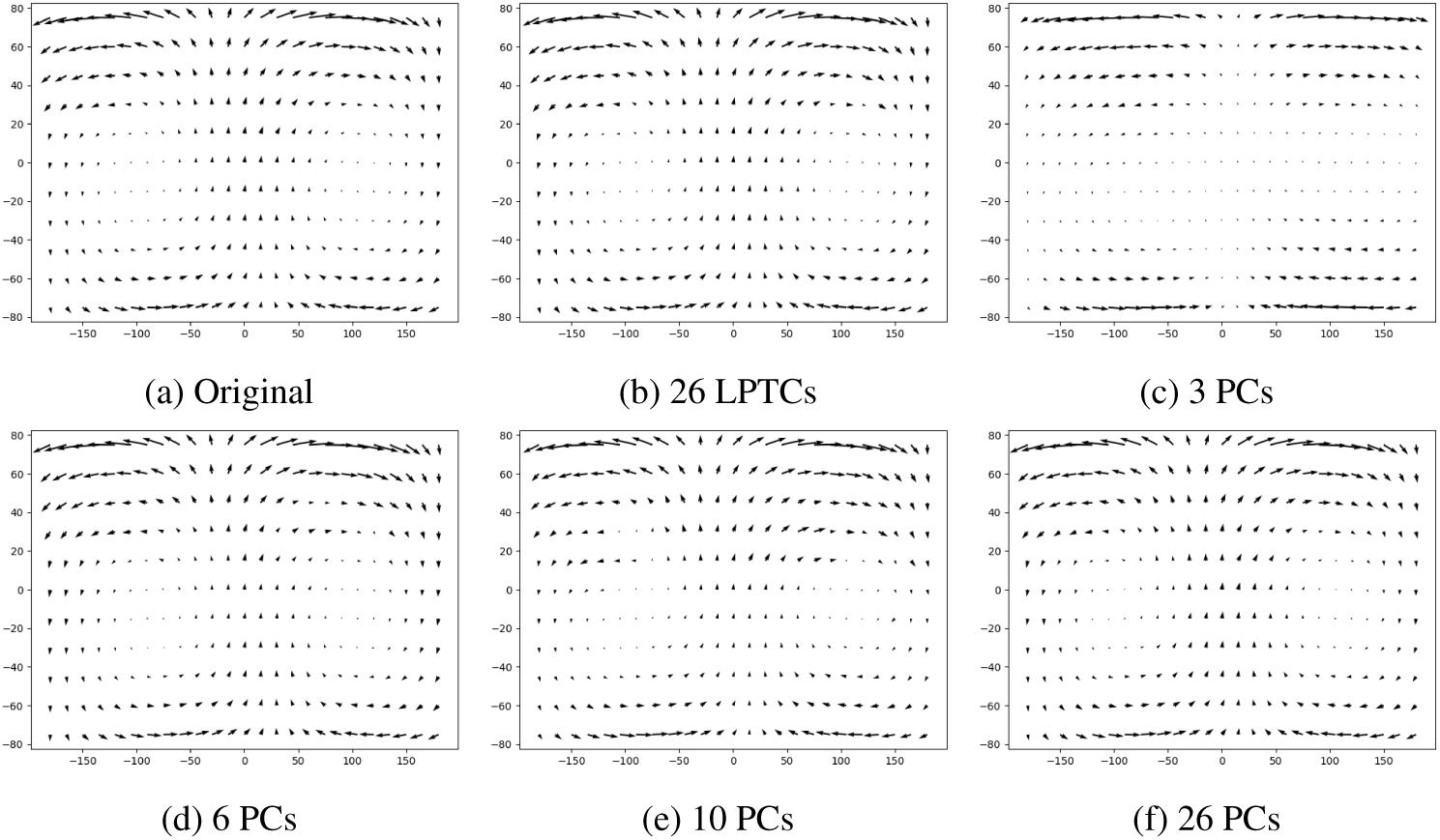
Example 1 of Optical Flow maps predicted. Original optical flow (a) encoded by 26 LPTCs(b), 3 PCs (c), 6 PCs (d), 10 PCs (e), and 26 PCs (f) and decoded back by the Gaussian Process model trained on 30% of the drone dataset.

**Figure 21:**
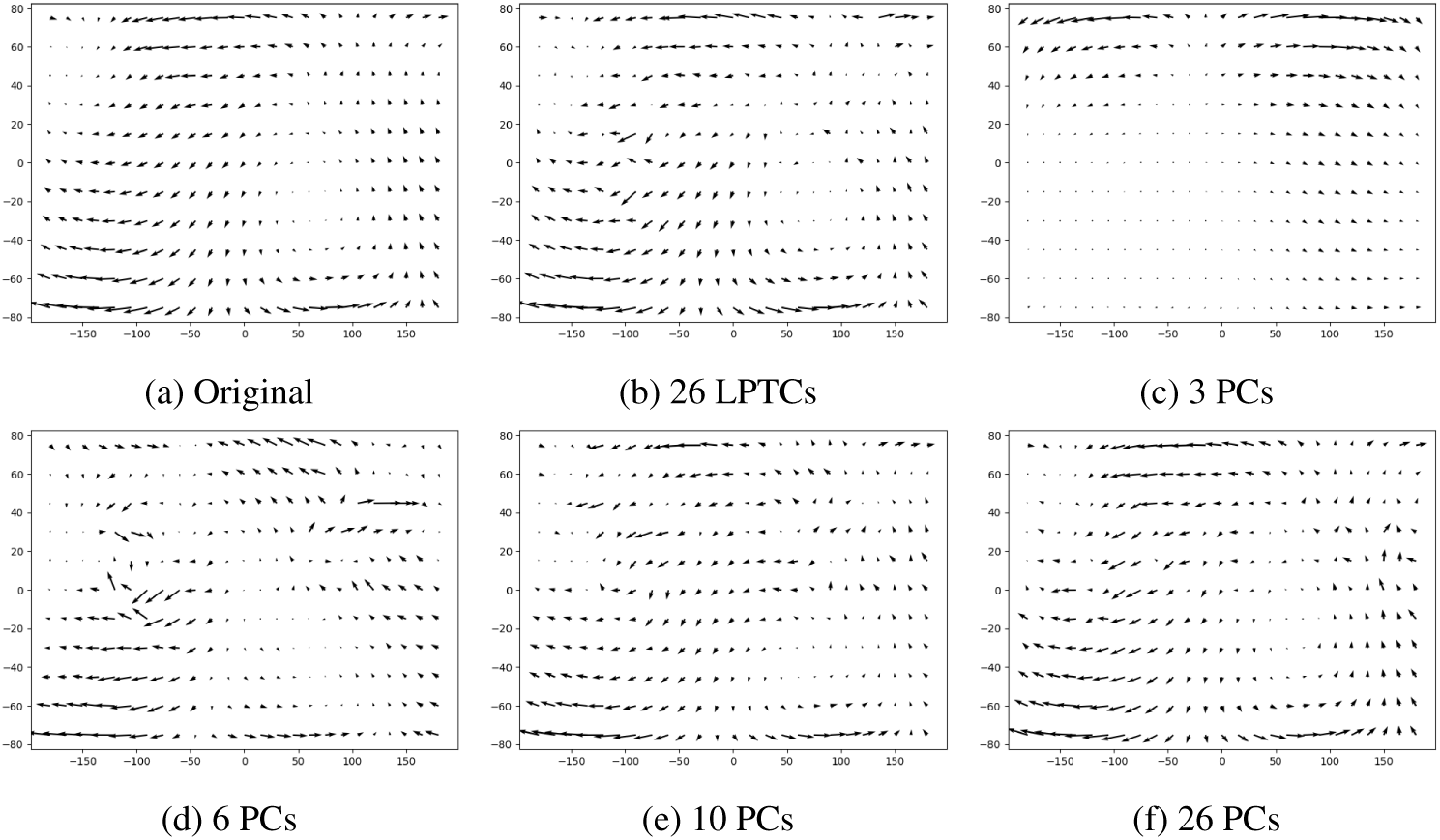
Example 2 of Optical Flow maps predicted. Original optical flow (a) encoded by 26 LPTCs(b), 3 PCs (c), 6 PCs (d), 10 PCs (e), and 26 PCs (f) and decoded back by the Gaussian Process model trained on 30% of the drone dataset.

**Figure 22:**
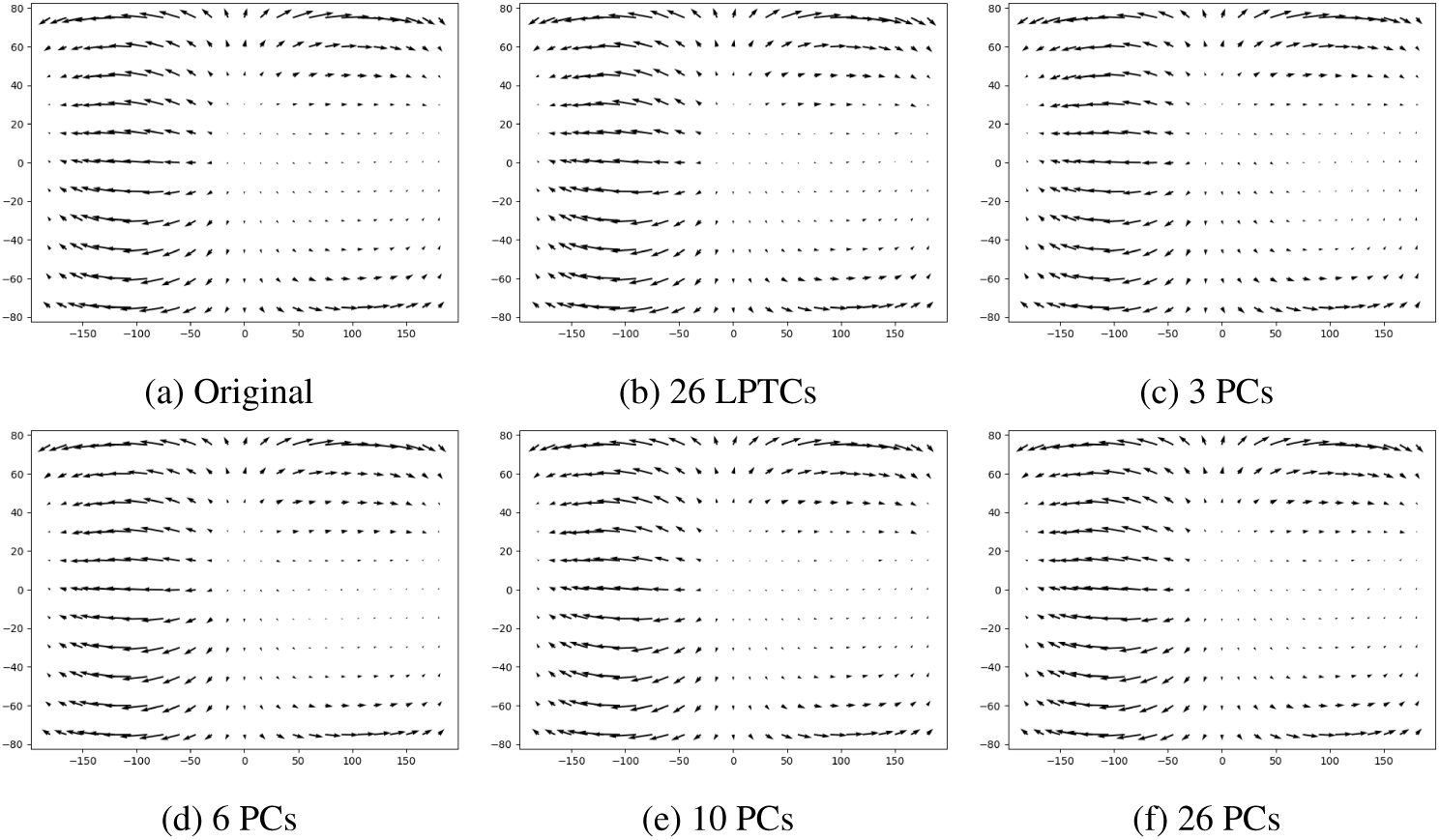
Example 3 of Optical Flow maps predicted. Original optical flow (a) encoded by 26 LPTCs(b), 3 PCs (c), 6 PCs (d), 10 PCs (e), and 26 PCs (f) and decoded back by the Gaussian Process model trained on 30% of the drone dataset.

**Figure 23:**
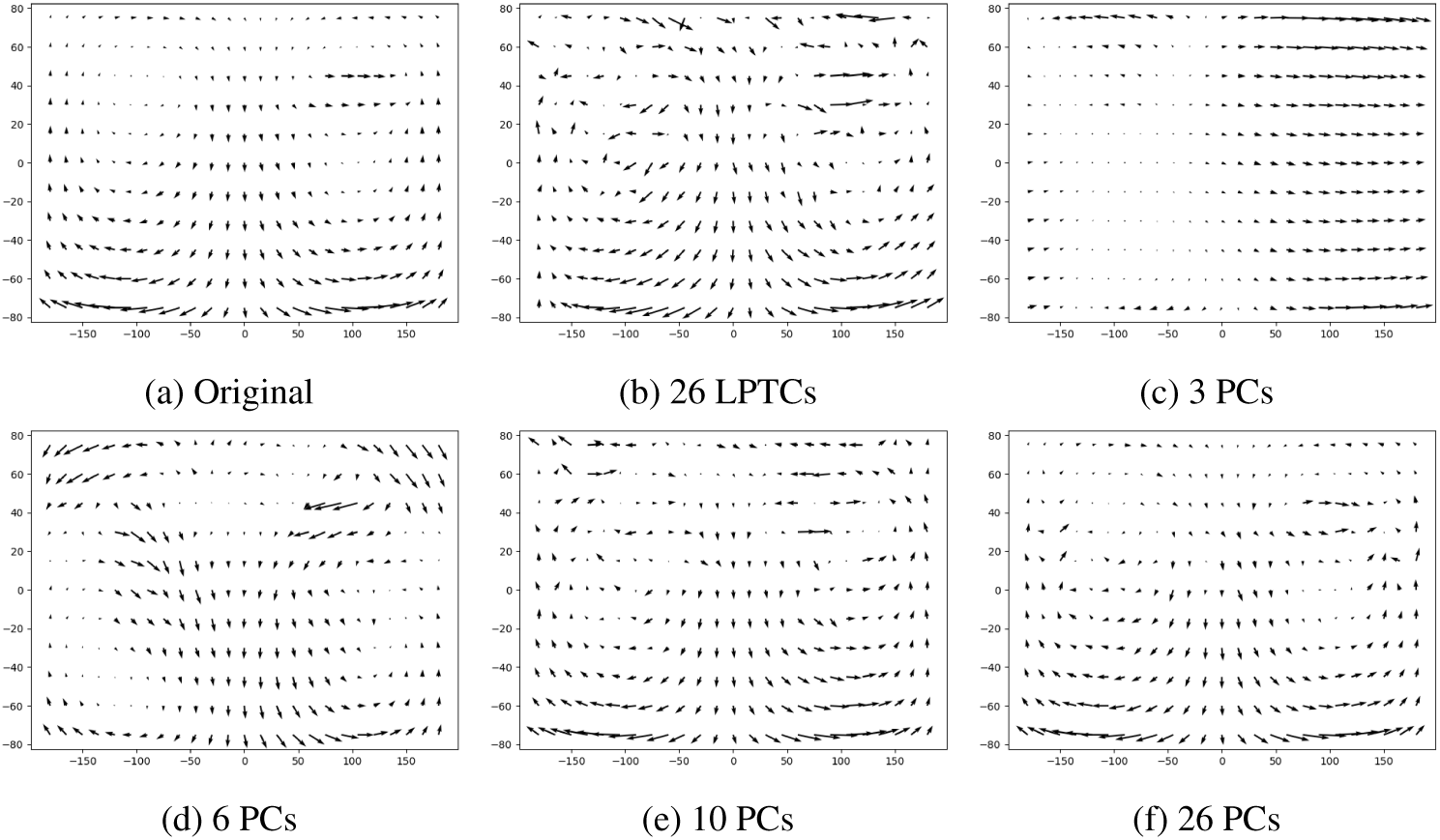
Example 4 of Optical Flow maps predicted. Original optical flow (a) encoded by 26 LPTCs(b), 3 PCs (c), 6 PCs (d), 10 PCs (e), and 26 PCs (f) and decoded back by the Gaussian Process model trained on 30% of the drone dataset. With this optical flow, the introduction of a discontinuity around 90 degrees azimuth and 40 degrees elevation generated by a close object the drone is fly near of leads to difficulty in estimating correctly the flow accurately and to numerous artefacts within the estimated flow maps.

**Figure 24:**
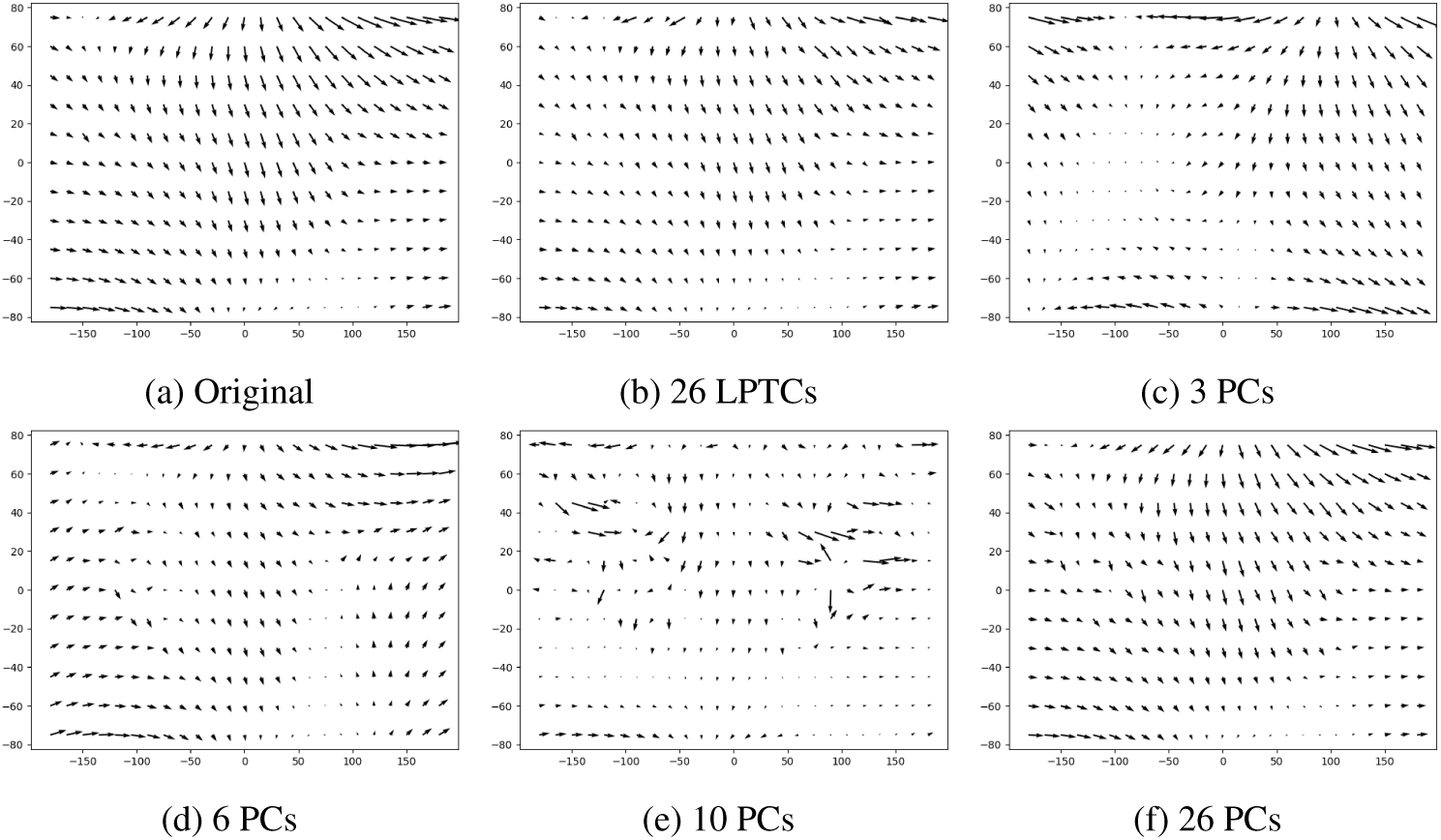
Example 5 of Optical Flow maps predicted. Original optical flow (a) encoded by 26 LPTCs(b), 3 PCs (c), 6 PCs (d), 10 PCs (e), and 26 PCs (f) and decoded back by the Gaussian Process model trained on 30% of the drone dataset. With such a motion, 10 PCs struggled at capturing the global flow with numerous outliers.

**Figure 25:**
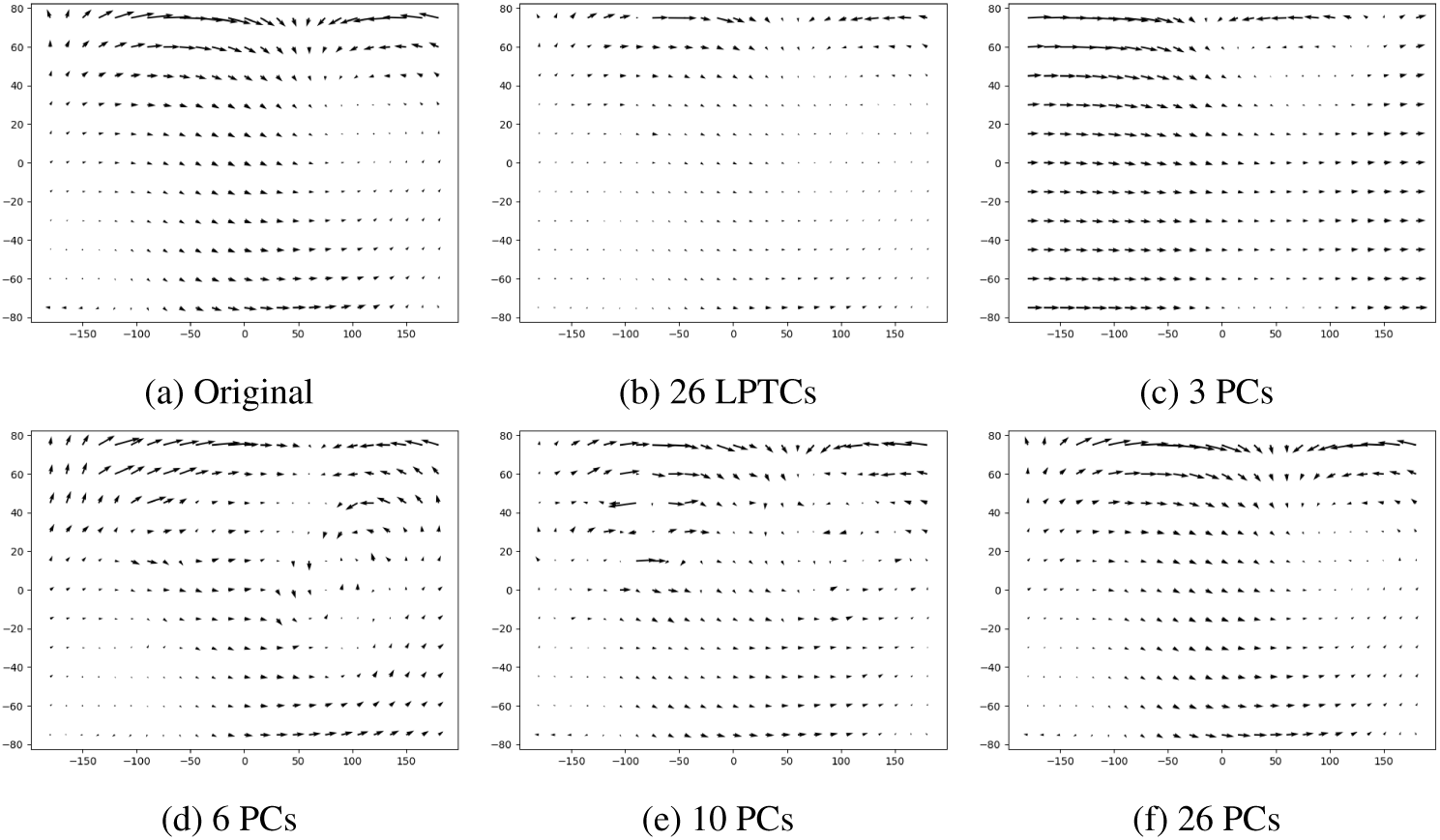
Example 6 of Optical Flow maps predicted. Original optical flow (a) encoded by 26 LPTCs(b), 3 PCs (c), 6 PCs (d), 10 PCs (e), and 26 PCs (f) and decoded back by the Gaussian Process model trained on 30% of the drone dataset.

**Figure 26:**
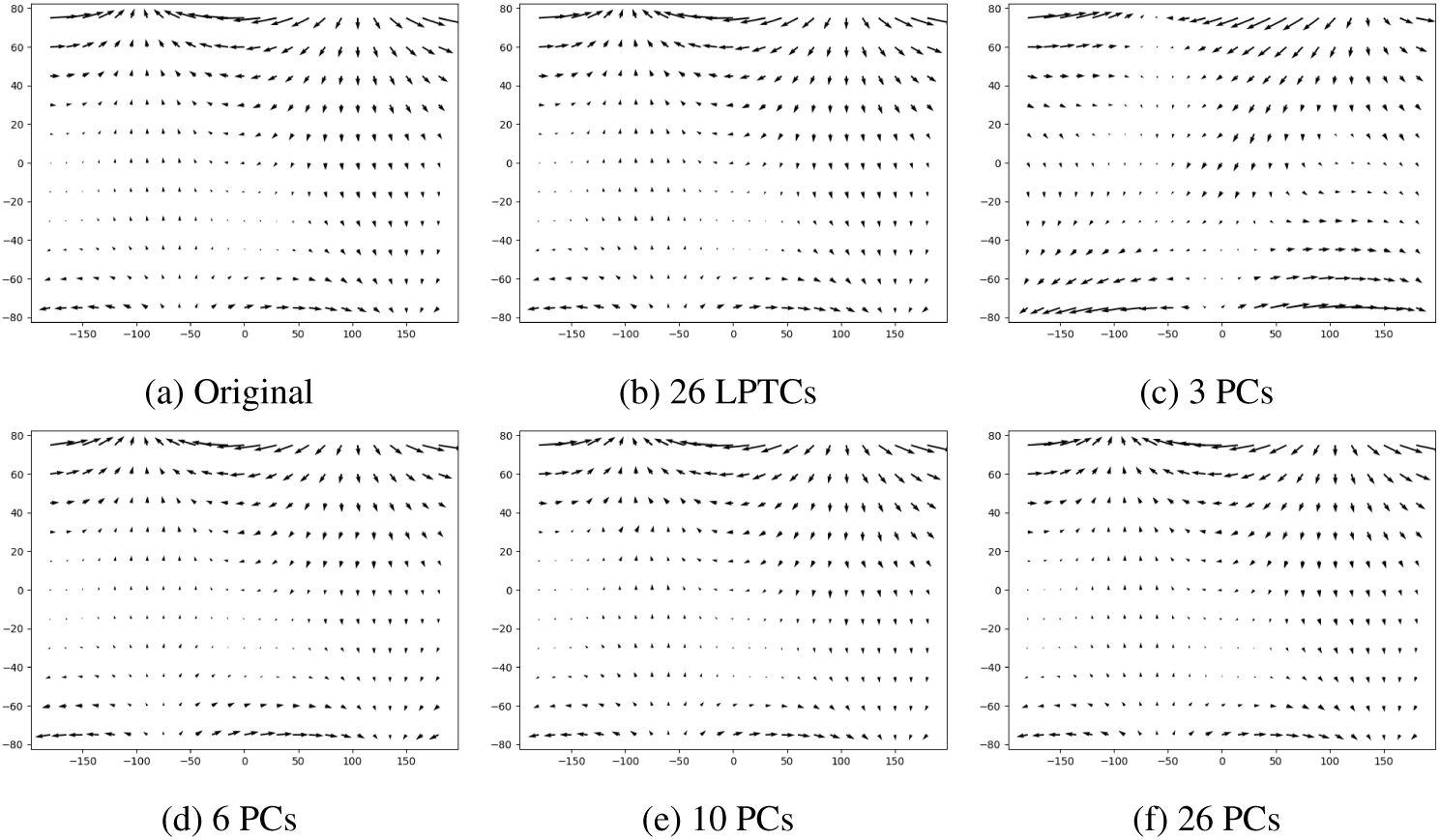
Example 7 of Optical Flow maps predicted. Original optical flow (a) encoded by 26 LPTCs(b), 3 PCs (c), 6 PCs (d), 10 PCs (e), and 26 PCs (f) and decoded back by the Gaussian Process model trained on 30% of the drone dataset.

